# Bacterial pathogens deliver water/solute-permeable channels as a virulence strategy

**DOI:** 10.1101/2023.07.29.547699

**Authors:** Kinya Nomura, Felipe Andreazza, Jie Cheng, Ke Dong, Pei Zhou, Sheng Yang He

## Abstract

Many animal and plant pathogenic bacteria utilize a type III secretion system to deliver effector proteins into the host cell^1,2^. Elucidation of how these effector proteins function in the host cell is critical for understanding infectious diseases in animals and plants^3–5^. The widely conserved AvrE/DspE-family effectors play a central role in the pathogenesis of diverse phytopathogenic bacteria^6^. These conserved effectors are involved in the induction of “water-soaking” and host cell death that are conducive to bacterial multiplication in infected tissues. However, the exact biochemical functions of AvrE/DspE-family effectors have been recalcitrant to mechanistic understanding for three decades. Here we show that AvrE/DspE-family effectors fold into a β-barrel structure that resembles bacterial porins. Expression of AvrE and DspE in *Xenopus* oocytes results in (i) inward and outward currents, (ii) permeability to water and (iii) osmolarity-dependent oocyte swelling and bursting. Liposome reconstitution confirmed that the DspE channel alone is sufficient to allow the passage of small molecules such as fluorescein dye. Targeted screening of chemical blockers based on the predicted pore size (15-20 Å) of the DspE channel identified polyamidoamine (PAMAM) dendrimers as inhibitors of the DspE/AvrE channels. Remarkably, PAMAMs broadly inhibit AvrE/DspE virulence activities in *Xenopus* oocytes and during *Erwinia amylovora* and *Pseudomonas syringae* infections. Thus, we have unraveled the enigmatic function of a centrally important family of bacterial effectors with significant conceptual and practical implications in the study of bacterial pathogenesis.

---

All AvrE/DspE-family effectors examined, including AvrE from *Pseudomonas syringae*, WtsE from *Pantoea stewartii*, DspA/E (DspE hereinafter) from *Erwinia amylovora*, and DspE from *Pectobacterium carotovorum*, are major virulence factors responsible for bacterial multiplication and induction of major disease symptoms including water-soaking and host cell death during infection^6–19^. AvrE/DspE-family effectors have been very challenging to study due to their extremely large size (approximately 200 kDa), high toxicity to plant and yeast cells, and sharing little sequence similarities to proteins of known function^20,21^. Several AvrE/DspE-family effectors were reported to interact with several plant proteins, including plant protein phosphatase PP2A subunits, type one protein phosphatases (TOPPs) and receptor-like kinases^22–25^. In addition, a yeast *cdc55* mutation affecting Cdc55-PP2A protein phosphatase activity was found to suppress DspA/E-induced yeast growth arrest^21^. While these interactions associate AvrE/DspE-family effectors to various host cellular processes, the fundamental question regarding the actual biochemical function of AvrE/DspE-family effectors has remained elusive.

In this study, we performed AlphaFold2 analysis of the 3-D models of AvrE/DspE-family proteins. Unexpectedly, AlphaFold2 predicts that this family of proteins fold into a porin-like β-barrel structure. This surprising prediction prompted us to conduct a series of cryo-EM imaging-, *Xenopus* oocyte-, liposome-, and *in planta* experiments. Our results show that AvrE/DspE-family effectors are water/solute-permeable channels that can be blocked by polyamidoamine (PAMAM) dendrimers. The unexpected discovery of the water/solute-permeable channel function of AvrE/DspE-family effectors solves a decades-long puzzle regarding one of the most important families of phytobacterial type III effectors and marks a major advancement in understanding bacterial pathogenesis.

## AlphaFold2 predication and cryo-EM imaging

To gain functional insights into the AvrE/DspE family of bacterial effectors, we constructed their predicted 3D models using the recently published AlphaFold2^26^ based on the fast homology search of MMseqs2^27^. The predicted AlphaFold2 models of DspE from *E. amylovora,* DspE from *P. carotovorum,* AvrE from *P. s.* pv. *tomato* (*Pst*) DC3000 and WtsE from *P. stewartia* (Fig. 1 and Extended Data Fig. 1 & 2) all reveal an overall similar architecture resembling a mushroom, with a prominent central β-barrel forming the stem, which is surrounded by a globular N-terminal domain (*E. amylovora* DspE: K298-H672), a WD40 repeat domain (H673-P912), and two perpendicularly arranged helix bundles (E998-T1222 and A1567-H1647) on the top. The predicted domain arrangement is supported by our cryo-EM imaging of *E. amylovora* DspE, where the 2D class averages clearly reveal an overall similar top view as the AlphaFold model, with circularly arranged globular domains surrounding a central pore (Fig. 1a,b).

**Figure 1.**
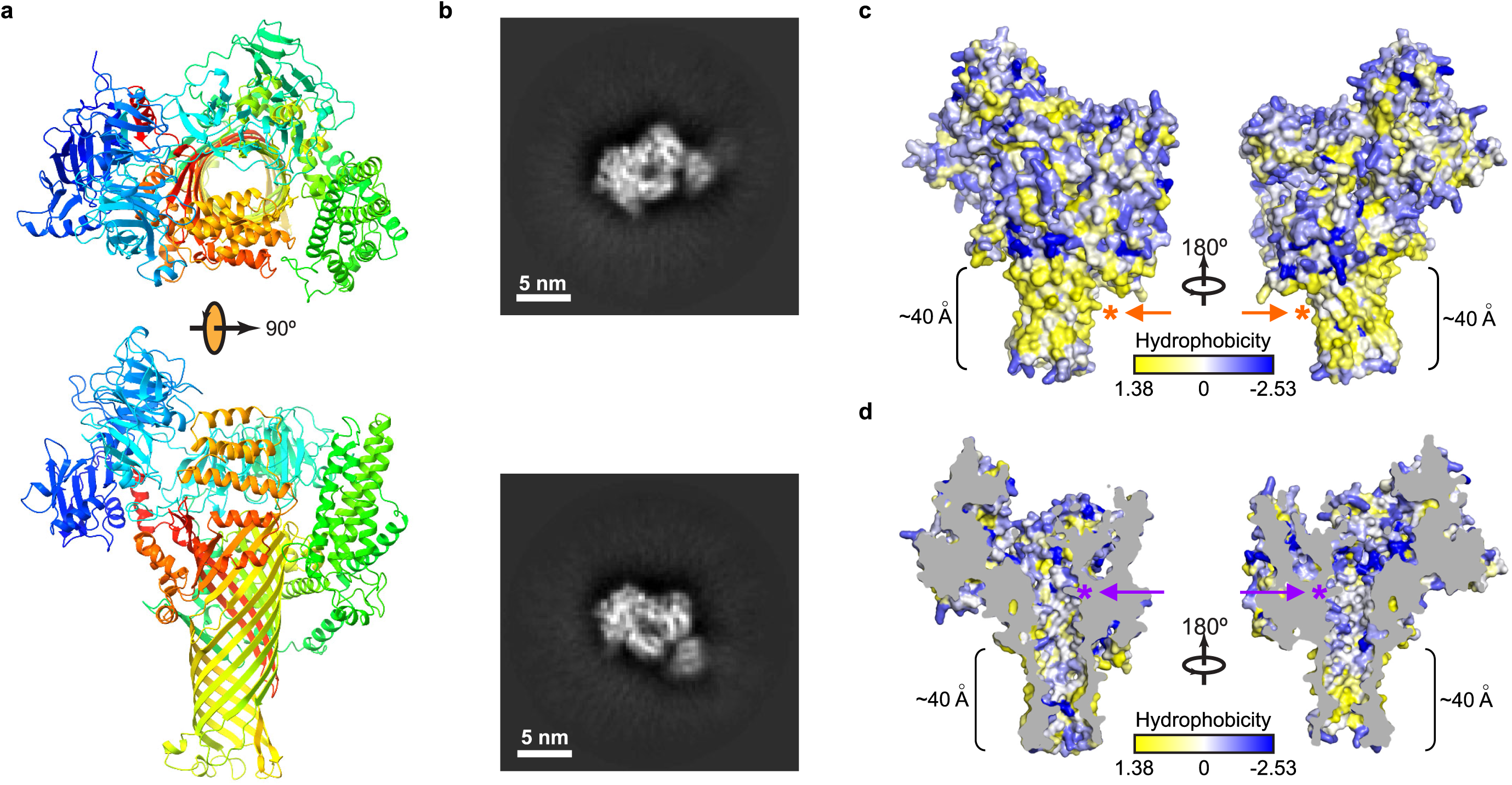
Model and cryo-EM images of *E. amylovora* DspE. **a,** 3D model of *E. amylovora* DspE generated by AlphaFold2 using MMseqs2. DspE (residue 298-1838) is shown in the cartoon model in rainbow colour, with N-terminus coloured in blue and C-terminus coloured in red. **b**, Cryo-EM 2D class averages of DspE, revealing a circular arrangement of domains around a pore. **c**, Surface representation of DspE. **d**, Sliced view of DspE. In panels c and d, residues are coloured based on their hydrophobicity scale. The length of the proposed membrane-spanning β-barrel stem is labeled. Orange and purple asterisks mark the approximate locations of the L1776/L1777/L1778 hydrophobic cluster and K1399/K1401 basic cluster, respectively.

Further examination of the predicted *E. amylovora* DspE 3D model reveals that surface β-barrel residues facing outside are enriched with hydrophobic amino acids (Fig. 1c), whereas inward-facing pore residues are predominantly hydrophilic (Fig. 1d). The length of the lower portion of the β-barrel stem covered by hydrophobic residues is estimated to be ∼40 Å, roughly the thickness of a cellular membrane. As AvrE has previously been reported to be membrane anchored^20^, the β-barrel stem of AvrE/DspE-family effectors likely inserts into the membrane and functions as a channel, similar to that of bacterial porins^28^. Such a mode of insertion is distinct from pore-forming bacterial toxins, such as *staphylococcal* alpha-hemolysin and *Clostridium perfringens* β-toxin, where the β-barrel is assembled through oligomerization of two long β-strands^29,30^.

## Expression of DspE and AvrE generates ionic currents in *Xenopus* oocytes

The unexpected prediction that AvrE/DspE-family effectors are channel-forming proteins prompted us to conduct experiments to test the hypothesis that AvrE and DspE may allow ion conductance when expressed in *Xenopus* oocytes. As shown in the current-voltage relation (Fig. 2b,c), inward currents and outward currents at negative and positive test potentials, respectively, were detected from oocytes injected with *dspE* or *avrE* cRNA. The reversal potentials for DspE and AvrE channels are ca. −25 mV. The DspE currents were not affected by niflumic acid, which blocks an endogenous Ca^2+^-activated chloride channels^31^, or fipronil, which inhibits GABA-gated Cl^−^ channels and glutamate-gated Cl^−^ channels^32^ (Extended Data Fig. 3a,b). Surface biotinylation experiment with oocytes expressing DspE confirmed that this protein is anchored across the oocyte membrane (Extended Data Fig. 4a).

**Figure 2.**
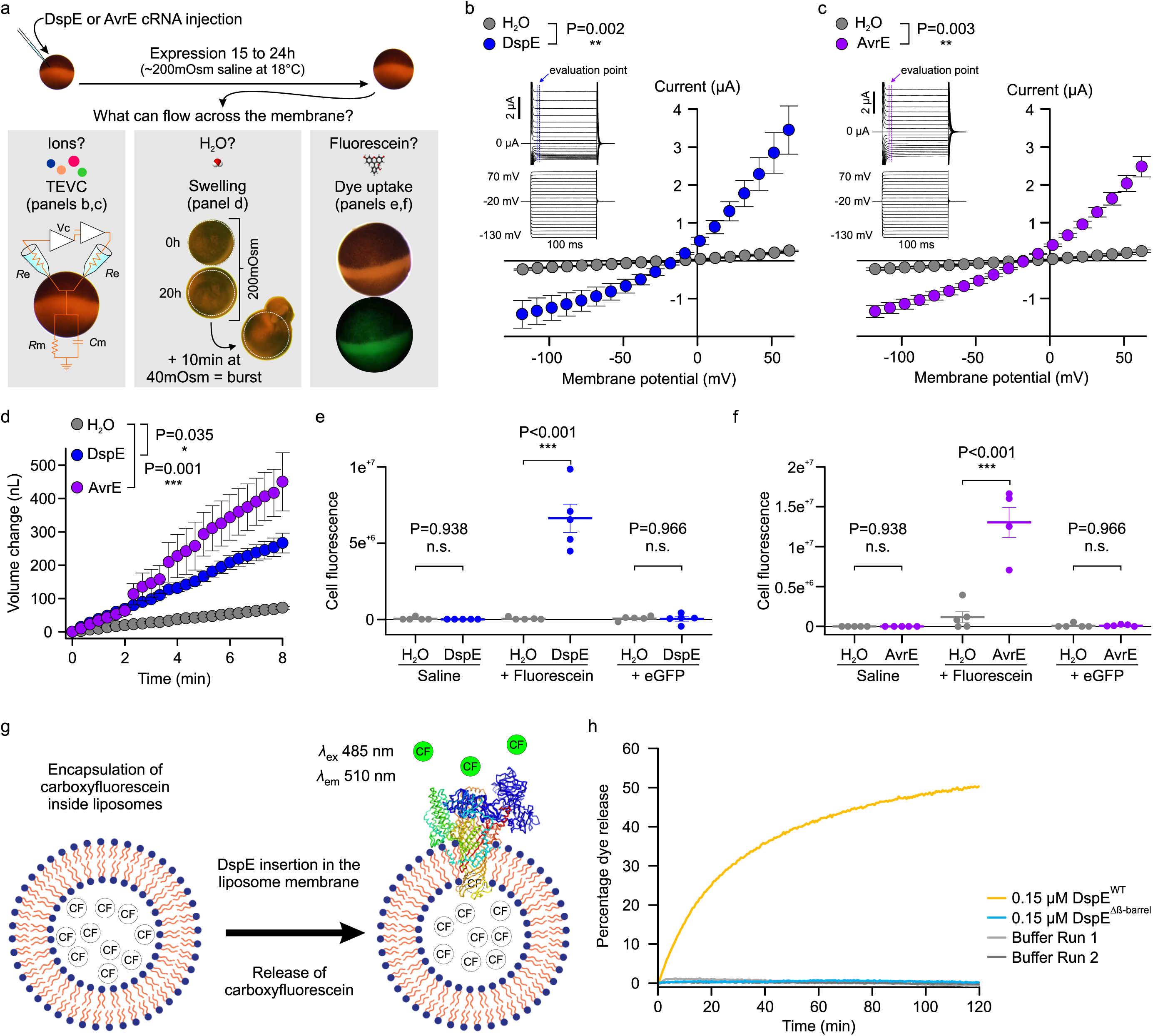
DspE and AvrE activities in *Xenopus* oocytes and liposome. **a,** Schematic of the three oocyte assays. Left: two-electrode voltage clamp (TEVC) to test ion conductance. Middle: Baseline and induced swelling/burst assay to test water conductance. Right: Dye uptake to test conductance to molecules larger than single ions. **b,c,** DspE and AvrE induce ion currents in TEVC assay. Mean ± SEM (n=5) current values at different test pulses from oocytes expressing DspE (0.01 ng cRNA/oocyte) or AvrE (0.1 ng cRNA/oocyte) were recorded. **d,** DspE (2 ng) and AvrE (20 ng) induced fast oocyte swelling/burst at 24 h after cRNA injection when placed in a low osmolarity (40 mOsm) solution. Mean ± SEM (n=5) values of increased oocyte volume in relation to its initial volume. **e,f,** Fluorescein/eGFP entry assays. Oocytes injected with 2 ng *dspE* (e) or 20 ng *avrE* (f) cRNA, or injected with water were incubated for 20 h in bath saline with or without fluorescein or eGFP. Values of fluorescence intensity were subtracted from the background and are present as mean ± SEM (n=5) corrected “total cell fluorescence”. **g,** Schematic illustration of DspE-dependent release of carboxylfluorescein (CF) encapsulated within a liposome. **h**, Fluorescence increased over time for CF-loaded liposome after addition of DspE (blue) or buffer (grey). The result is a representative of 3 experimental replicates. Two-way ANOVA (b,c,e,f) or two-way repeated measure ANOVA (d) values and exact P-values for all comparisons are detailed in the Source Data files.

We further characterized whole-cell currents from DspE-expressing oocytes by conducting ion permeability experiments. Replacing extracellular sodium in ND96 recording solution (see Methods) with potassium or other cations caused only minor variations on the magnitudes of DspE currents (Extended Data Fig. 3c). Similarly, only minor variations in the magnitudes of DspE currents were observed when extracellular chloride was replaced with various anions, except for Na-MES (Extended Data Fig. 3d,e). When 50% or 100% of the NaCl was replaced by Na-MES, progressively smaller outward currents were observed, and the reversal potential was shifted to a less negative value (Extended Data Fig. 3e). However, the negative reversal potential was not affected by replacement of other ions. These results suggest that Cl^−^ may play a major role in carrying the outward current and that DspE channel appears to have some selectivity toward anions including chloride. Future research is needed to comprehensively survey possible ion selectivity of the DspE channel.

## DspE and AvrE induce swelling and bursting of *Xenopus* oocytes

We noticed an interesting phenomenon during voltage-clamp current recording experiments: many oocytes injected with *dspE* or *avrE* cRNA showed baseline swelling (Extended Data Fig. 5a), reminiscent of oocytes expressing plant aquaporins^33^. This raised the possibility that AvrE and DspE proteins may function like aquaporin channels to allow water to pass through cell membranes along an osmotic gradient, assuming that the osmolarity of the oocyte bathing medium (∼200 mOsm) may be lower than that of the oocyte cytoplasm. To more directly test this possibility, we adopted an oocyte swelling assay used for aquaporins and transferred oocytes from 200 mOsm to 40 mOsm bathing medium to create a larger osmotic difference. Strikingly, both DspE- and AvrE-expressing oocytes dramatically swelled (Fig. 2d) and eventually burst (Supplementary Videos 1 and 2). We conducted further experiments to determine if plant cells expressing AvrE would swell using our previously produced transgenic DEX::*avrE* plants^20^. As shown in Extended Data Fig. 5b,c, Arabidopsis leaf protoplasts expressing AvrE swelled to a greater extent compared to control Arabidopsis leaf protoplasts which have endogenous aquaporins. This provides further evidence that the AvrE channel has an ability to increase water permeability *in planta*^7–19^.

## Size-dependent AvrE/DspE channel selectivity

The predicted AvrE/DspE channels have a diameter of 15-20 Å, significantly larger than the size of a water molecule or a simple ion. We hypothesized that, in addition to ions and water, the AvrE/DspE channels may allow larger molecules to pass through. We conducted fluorescent dye permeability assays to test if molecules smaller than the predicted pore size could pass through the membrane, whereas molecules larger than the predicted pore size could not. Two fluorescent molecules were tested: fluorescein (MW of 332 Da with an estimated maximum molecular diameter of 7 Å) and green fluorescent protein (eGFP) (MW of 27 kDa with an estimated minimum diameter of 30 Å). As shown in Fig. 2e, 2f, fluorescein entered oocytes expressing DspE or AvrE, whereas eGFP could not, consistent with the predicted AvrE/DspE channel diameter. Notably, liposome-based *in vitro* reconstitution of DspE was sufficient to cause time-dependent release of carboxyfluorescein (MW of 376 Da) encapsulated within soybean liposomes, whereas neither the buffer control nor the DspE^Δβ-barrel^ mutant, in which the majority of β-barrel-forming sequences are deleted, displayed significant activity (Fig. 2g,h), suggesting that no other protein is needed for baseline passage of small molecules through the DspE channel. Similar to the oocyte experiment, although DspE readily induced carboxyfluorescein dye release from liposomes (Fig. 2h), no time-dependent release of large molecules, such as FITC-polysucrose 40 (MW of 30-50 kDa with an estimated diameter of 80 Å), was observed (Extended Data Fig. 5d).

## Mutational analysis of the DspE channel

We made several mutant derivatives of DspE to evaluate their functional consequences. As a negative control, DspE^Δβ-barrel^ failed to induce water-soaking symptom in *N. benthamiana* leaves, to conduct ion currents, to cause oocyte swelling, or to allow fluorescein release in liposome assay (Fig. 2h; Extended Data Fig. 6). Similarly, triple mutation of three conserved hydrophobic, outward-facing residues (L1776, L1777, L1778) of the predicted transmembrane region of DspE (location indicated by the orange asterisk in Fig. 1c) also abolished the DspE activities (Extended Data Fig. 6). Surface biotinylation of oocytes expressing DspE^Δβ-barrel^ and DspE^L1776E/L1777E/L1778E^ showed that they are no longer accessible by surface biotinylation and therefore likely cannot anchor across the membrane (Extended Data Fig. 4a). Finally, charge-reversal double mutations at K1399 and K1401, two inward-facing residues of the β-barrel, (location indicated by the purple asterisk in Fig. 1d, not conserved among AvrE-family members) partially abolished the DspE activities (Extended Data Fig. 6). Due to the novel nature of the DspE channel, future research is needed to comprehensively define inward-facing residues that are critical for the DspE function.

## PAMAMs G0 and G1 are inhibitors of DspE/AvrE channels

The discovery of AvrE and DspE functioning as channels offered an opportunity to identify compounds whose molecular diameters could fit the predicted pores of AvrE/DspE channels and might therefore block AvrE/DspE activities. We focused on a class of synthetic polyamidoamine (PAMAM) dendrimers, which have programmable molecular diameters^34^. For example, PAMAM G0 has a diameter of 15 Å, whereas PAMAM G1 has a diameter of 22 Å (https://www.dendritech.com/pamam.html). We found that the currents passing through the DspE and AvrE channels were reduced by G1 in a dose-dependent manner, reaching 71% of inhibition on DspE channel and 93% of inhibition on AvrE channel, respectively, at 10 mM of G1 at 50 mV test pulse (Fig. 3a). Similar inhibition by G0 was observed on oocytes expressing DspE, reaching 68% inhibition at 10 mM G0 at 50 mV test pulse (Extended Data Fig. 7a).

**Figure 3.**
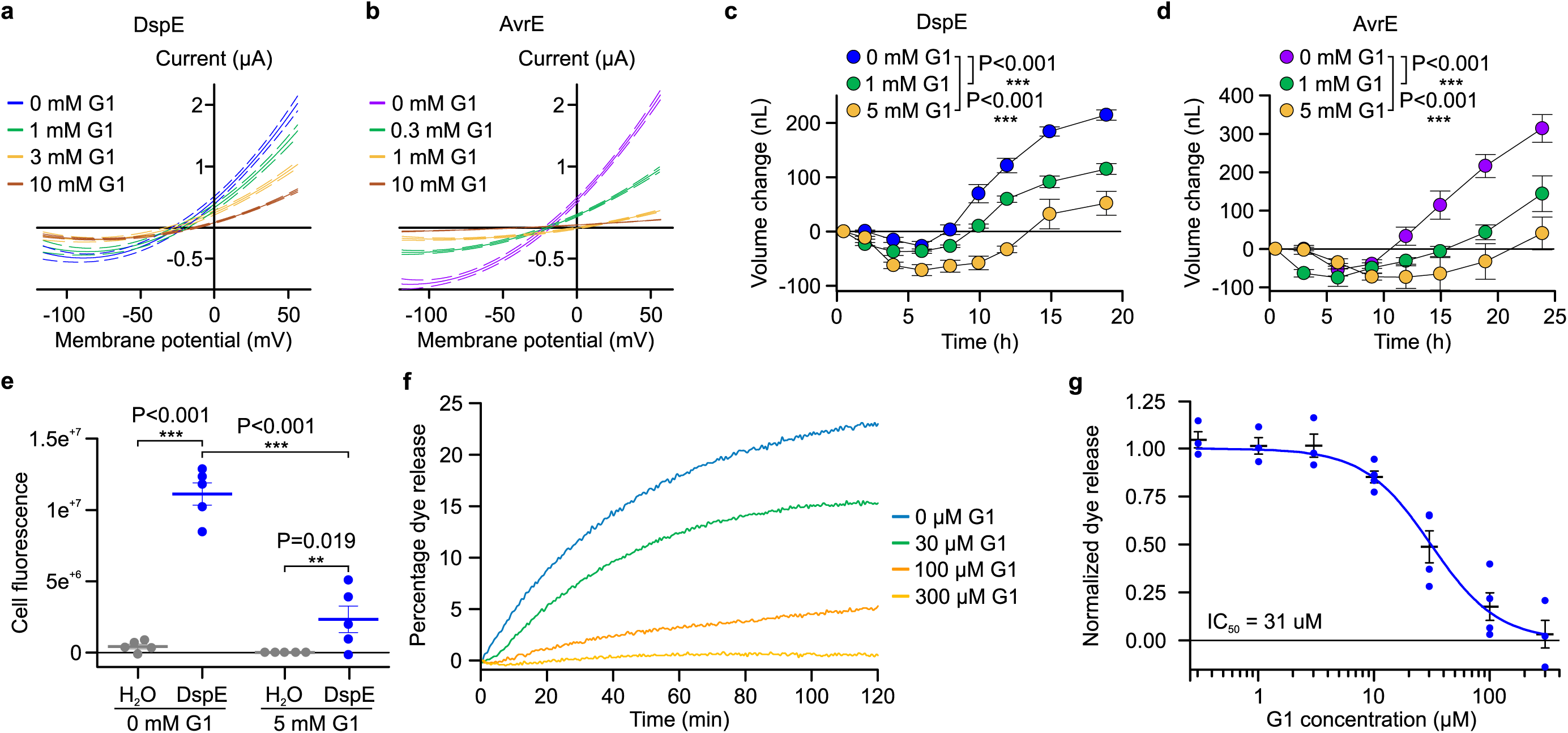
Inhibition of DspE and AvrE channels in oocytes by polyamidoamine (PAMAM) dendrimers G0 and G1. Assay flowchart followed Fig. 1a, except with inhibitors added to the bath saline. **a,b,** DspE- and AvrE-dependent currents were inhibited by G1. Solid lines represent fit values across the entire voltage range, with dashed lines representing the lower and upper 95% confidence interval of a quadratic polynomial regression for each treatment after subtracting control values. **c**,**d,** Baseline swelling of oocytes injected with 1 ng *dspE* (c) or 20 ng *avrE* (d) cRNA was reduced in the presence of G1. Data shows mean ± SEM (n=5) increase in volume from the start point (at the time of injection) after subtracting control values. **e,** Inhibition of fluorescein uptake. G1 reduced fluorescein entry into oocytes expressing DspE as evaluated 6 h after injection with 1 ng *dspE*/oocyte. Data is presented as in Fig. 2e. **f,** DspE-mediated dye release from liposome in the presence of increasing concentrations of G1. The result is a representative of 3 experimental replicates. **g,** Dose-dependent inhibition of the liposome dye release. IC_50_: half maximal inhibitor concentration. Data shows mean ± SEM (n=3 for 0.3, 1, 3, and 300 µM G1; n=4 for 10, 30, and 100 µM G1). Experiments were independently performed two times for a,b,e, four times for c,d, and seven times for f,g. Two-way ANOVA (e) or two-way repeated measure ANOVA (c,d) values and exact P-values for all comparisons are detailed in the Source Data files.

When G1 was added to the ND96 incubation buffer the baseline swelling over time was also inhibited in a dose-dependent manner, reaching a maximum of 76% of inhibition for 5 mM of G1 at 19 h after injection (Fig. 3c). Again, a similar effect was observed when this inhibitor was tested on AvrE-expressing oocytes, with 89% of inhibition for 5 mM of G1 (Fig. 3d).

We also tested the effect of G1 on fluorescein uptake by oocytes expressing DspE, and found it inhibited fluorescent dye uptake, reaching 79% inhibition by the end of the assay (Fig. 3e). Finally, we tested G1 on purified DspE protein reconstituted in liposomes using the DspE-dependent liposome dye release assay. We found that G1 dose-dependently inhibited the release of fluorescein from the soybean liposomes after the addition of DspE (Fig. 3f). Fitting of the dye release yielded an IC_50_ value of 31 µM for G1 (Fig. 3g). Taken together, we have successfully identified PAMAM G1 as an inhibitor of AvrE/DspE-family channels.

## PAMAM G1 inhibits bacterial infection

The ability of PAMAM G1 to inhibit DspE and AvrE activities in oocytes and liposomes *in vitro* raised the exciting possibility that we had identified a lead compound that could interfere with the AvrE/DspE virulence function *in planta* during bacterial infection. We first tested this possibility against the AvrE function during *P. syringae* infection. In many *P. syringae* strains, AvrE is functionally redundant to another effector, HopM1^11,13,19^. Mutation of either *avrE* or *hopM1* alone does not strongly affect *Pst* DC3000 virulence, but the *avrE hopM1* double mutant is severely impaired in virulence^11,19^. Interestingly, while *avrE*-family effector genes are conserved widely^35^, *hopM1*-famly genes and/or their secretion chaperon genes are subjected to natural genetic mutations as in the case of the major pandemic bacterial pathogen *Pseudomonas syringae* pv. *actinidiae*^36^. We found that G1 effectively inhibited *Pst* DC3000 infection of Arabidopsis in an AvrE-dependent manner (i.e., inhibition occurs in the *hopM1* deletion mutant, which simulates natural mutations in the *hopM1* gene; Fig. 4a and 4b). Furthermore, inhibition of AvrE function by PAMAM G1 was not associated with induction of the PR1 protein, a marker for activation of salicylic acid-dependent immune responses in plants (Fig. 4c), or with negative effects on plant appearance (Extended Data Fig. 7b) or seed production (Extended Data Fig. 7c). Next, we tested PAMAM G1 against *Erwinia amylovora* infection. In *E. amylovora*, DspE plays an essential role in causing the devastating fire blight diseases, as the *dspA/E* mutant is largely nonpathogenic^7,8^. Remarkably, we found that G1 completely inhibited *E. amylovora* infection of highly susceptible pear fruits, phenocopying the *dspE* mutant of *E. amylovora*, an observation consistent with DspE being an indispensable virulence effector (Fig. 4d).

**Figure 4:**
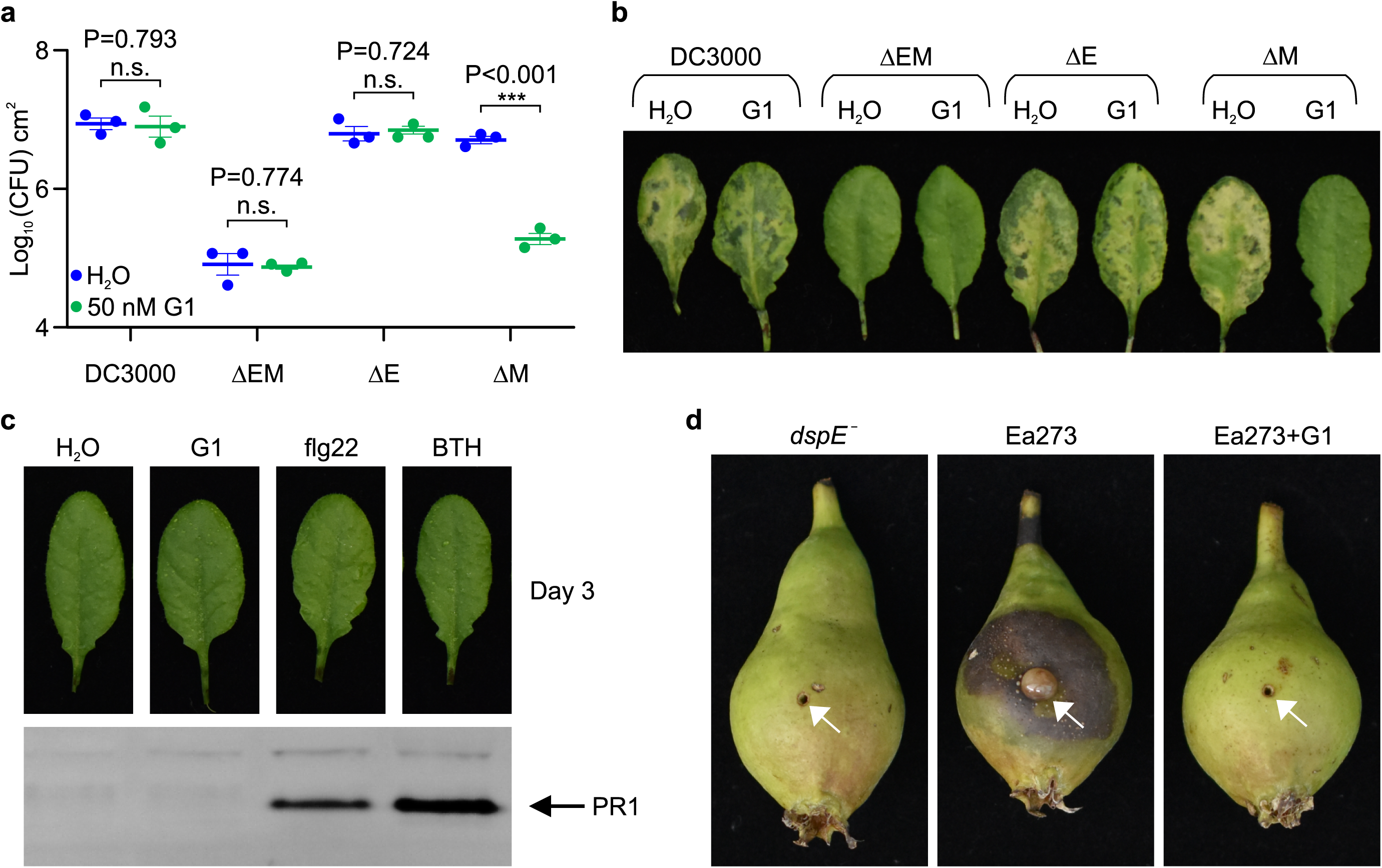
Effect of PAMAM G1 on bacterial infections. **a,b,** PAMAM G1 inhibits *Pseudomonas syringae* pv. *tomato* (*Pst*) DC3000 multiplication in an AvrE-dependent manner. 1×10^6^ CFU/mL of *Pst* DC3000, the *avrE hopM1* double mutant (ΔEM), the *avrE* single mutant (ΔE) or the *hopM1* single mutant (ΔM) were syringe-inoculated into leaves of Arabidopsis wild-type Col-0 plants, with or without 50 nM PAMAM G1. Bacteria populations (mean ± SEM; n=3) (a) in leaves were determined at day 3 post infiltration. Disease symptom picture (b) was taken at day 4 post infiltration. **c,** PAMAM G1 does not induce PR1 protein expression in Arabidopsis. Arabidopsis Col-0 leaves were syringe-infiltrated with 10 µM PAMAM G1. Plants were kept under high humidity (>95%) for 3 days at 23L. PR1 protein in leaves was detected by an α-PR1 polyclonal antibody. 1 µM flg22 and 100 µM BTH are inducers of PR1 expression. Gel image cropping is shown in Supplementary Information Fig. 2. **d,** PAMAM G1 inhibits fire blight disease by *Erwinia amylovora* Ea273. Immature pear fruits were spot-inoculated (indicated by arrows) with 10 µL of 1×10^3^ CFU/mL of Ea273 or the *dspE* mutant, with or without 10 µM PAMAM G1. Inoculated pears were placed on a wet paper towel in sterile box and incubate at 28L for 10 days. Diseased pears show a dark, necrotic appearance. Experiments were performed three times with similar results. Two-way ANOVA (a) values and exact P-values for all comparisons are detailed in the Source Data files.

It is well known that type III secretion systems and their effectors are only expressed and needed for bacterial growth in host tissues, but not *in vitro*^1,2^. In accordance with this, G1 did not inhibit *E. amylovora* or *Pst* DC3000 growth *in vitro* (Extended Data Fig. 7d), providing further evidence that PAMAM G1 is truly a AvrE/DspE-specific virulence inhibitor, not just a nonspecific bactericidal antibiotic.

## DISCUSSION

Since the initial report of AvrE in *P. syringae* almost three decades ago^37^, the central importance of AvrE/DspE-family effectors in diverse pathogenic bacterial species has attracted the attention of researchers. Prior to this study, however, researchers largely pursued a working hypothesis that, like most other type III effectors, AvrE/DspE-family effectors would target host proteins to exert their virulence functions. Our finding that AvrE/DspE-family effectors could directly function as water/solute-permeable channels in this study is therefore striking and unexpected. We propose a new integrated model for the function of AvrE/DspE-family effectors (Extended Data Fig. 8). The C-terminal halves of AvrE/DspE-family effectors act primarily as a novel class of water/solute-permeable channels dedicated to creating osmotic/water potential perturbation and a water/nutrient-rich apoplast in which bacteria multiply within the infected plant tissues. Future research is needed to determine how AvrE/DspE-family channel activity, discovered in this study, mediates water/solute/nutrient flows and apoplast osmolarity *in planta* to generate macroscopic water-soaking, host cell death and defense suppression in the infected tissue, as shown previously^6,13,19,25,38,39^. We found that infiltrating the Arabidopsis leaf apoplast with water (to simulate water-soaking) was sufficient to suppress flg22-induced callose deposition (Extended Data Fig. 9a,b), suggesting that water-soaking and suppression of certain immune responses (e.g., defense-associated callose deposition) are linked processes.

As large proteins with potentially many protein-interacting interfaces, AvrE/DspE-family effectors can additionally engage host proteins, including plant protein phosphatase PP2A subunits, type one protein phosphatases (TOPPs) and receptor-like kinases^22–25^, to impact aspects of AvrE/DspE functions. It is striking that the AvrE/DspE channel inhibitor PAMAM G1 can essentially phenocopy the avrE/dspE genetic mutations in abrogating all major virulence phenotypes associated with AvrE and DspE, including water-soaking, tissue necrosis as well as bacterial multiplication in infected tissues (Fig. 4). Therefore, future research should examine the possibility that some of the identified AvrE/DspE/WtsE-interacting host proteins may act through modulating AvrE-family channel properties and/or optimizing downstream pathogenic outcomes of AvrE-family channel activities.

To our knowledge this is the first time that bacterial type III effectors have been shown to directly function as pathogenic water/solute-permeable channels as a virulence mechanism. In addition to unraveling the long-sought-after function of AvrE/DspE-family effectors, this study identified a chemical inhibitor of AvrE/DspE channels that appears to be broadly effective in reducing AvrE/DspE-mediated bacterial infections. As such, the discovery of the water/solute-permeable channel function of AvrE/DspE-family effectors has broad implications in the study of bacterial pathogenesis and bacterial disease control in plants.

## Supporting information

Source Data for Extended Data Figures

Source Data for Extended Data Figure 2

Source Data for Extended Data Figure 3

Source Data for Extended Data Figure 4

## METHODS

### Cloning, expression and purification of *E. amylovora* DspE *and* DspE^Δβ-barrel^ in *E. coli*

The *dspE* gene was PCR-amplified from the genomic DNA of *Erwinia amylovora* strain Ea273 and cloned into a modified *pET28a* vector (MilliporeSigma) as C-terminal fusion to maltose-binding protein (MBP) containing a His_8_ tag, a preScission protease cleavage site (PPX), and a FLAG tag in the form of MBP-His_8_-PPX-FLAG-DspE (herein after as MBP-DspE). Plasmid-transformed BL21(DE3) *E. coli* cells were grown in Luria-Bertani (LB) medium at 37 °C until OD_600nm_ reached 0.4-0.6, and were then induced with 0.1 mM IPTG and grown at 18 °C overnight. Harvested cell pellets were resuspended in a lysis buffer containing 20 mM HEPES (pH 7.5), 300 mM NaCl, and 2.5% glycerol, supplemented with cOmplete EDTA-free protease inhibitor tablet (Roche) and DNase I, and lysed by French press. Following centrifugation at 20,000 rpm for 30 min at 4 °C to remove cell debris, the fusion protein was purified using TALON® Cobalt resin (Takara Bio). Following extensive washing in a buffer containing 20 mM HEPES (pH 7.5), 300 mM KCl, 2.5% glycerol, 1 mM ATP, 5 mM MgCl_2_, and 10 mM imidazole, the fusion protein was eluted in a buffer containing 20 mM HEPES (pH 7.5), 300 mM NaCl, 2.5% glycerol and 250 mM imidazole, and was further purified on a Superose^®^ 6 Increase 10/300 GL column (Cytiva Life Science) pre-equilibrated with a buffer containing 20 mM HEPES (pH 7.5), 150 mM NaCl and 1 mM DTT at 4 °C. The protein fractions at the peak were aliquoted and flash-frozen at −80 °C for storage.

The construct of DspE^Δβ-barrel^ (Δ1278-1566 + Δ1649-1813) was generated from the WT MBP-DspE construct described above through infusion cloning. The mutant protein was purified by following identical procedures as the WT protein.

Representative size exclusion chromatography profiles and SDS-PAGE gels of the purified MBP-DspE and MBP-DspE^Δβ-barrel^ proteins are shown in Extended Data Fig. 10a,b.

Constructs of MBP-DspE and MBP-DspE^Δβ-barrel^ were made by the infusion cloning methods using NEBuilder HiFi® DNA assembly kit (New England Biolabs) with the following primer sets:

WT DspE:

*dspE*_forward_primer: GATGGAATTAAAATCACTGGGAACTGAACACAAG

*dspE*_reverse_primer: GAAGGAAGGGCTGGAAATGAAGAGCTAATTGATTAA

Vector_forward_primer: GAGCTAATTGATTAATACCTAGGCTGCTAAACAAAG

Vector_reverse_primer: TTCTGTTCCAGGGGCCCGCGATGGAATTAAAATC

DspE^Δβ-barrel^ (Δ1278-1566 + Δ1649-1813):

*dspE* forward_primer: CCTGGACAGTGCGGAGCCGGTGACCAGCAA

*dspE* reverse_primer: AGGCTGCGGACAGCCACAGCGGAATAGCT

Vector_forward_primer: CACAGCGGAATAGCTCAGGCTAATCCGCAG

Vector_reverse_primer: GAATACGCTGTTGTCCCTGGACAGTGCGGA

### Cryo-EM sample preparation and data collection and processing

Cryo-EM grids were prepared using a Leica EM GP2 Automatic Plunge Freezer in a humidity-controlled chamber operated at 10 °C and 85-90% relative humidity. Homemade Gold Quantifoil^®^ R1.2/1.3 300-mesh grids were glow-discharged using the PELCO easiGlow^TM^ glow discharge cleaning system (TED PELLA, INC) before sample application. During sample freezing, a 3 μL sample of DspE (∼1.1 mg/mL) was applied to freshly glow-discharged grids and incubated on grids for 60 s before blotting with Whatman #1 filter paper (Whatman International Ltd) for 2.8 s. The grids were then immediately plunge-frozen in liquid ethane and stored in liquid nitrogen before data acquisition.

A total of 7,810 movies were recorded on an FEI Titan Krios electron microscope (Thermo Fisher Scientific) operated at 300 kV equipped with a K3 direct electron detector (Gatan, Inc.) operated in the counting mode. Movies were collected at a nominal magnification of 81,000x using pixel size of 1.08 Å/pix with a defocus range from −2.4 μm to −0.8 μm using Latitude^TM^ S (Version 3.51.3719.0, Gatan, Inc.) automated image acquisition package. Each stack was exposed for 2.8 s with an exposure time of 0.047 s per frame, resulting in 60 frames per stack. The total dose was approximately 56.3 e^−^/Å^2^ for each stack.

Movie alignment and contrast transfer function (CTF) estimation were performed with the patch motion correction model and patch CTF estimation module in cryoSPARC^40^. A total of 7,141 micrographs were selected from a total of 7,810 images based on the CTF fitting resolution using a cutoff value of 4.0 Å. A total of ∼2.4 M particles were picked using pre-trained TOPAZ^41^ models, of which ∼167 K particles corresponding to full-length protein with high resolution features were selected to generate 2D class averages. Cryo-EM samples of DspE so far showed a severe orientation bias of the particles, which prevented high-resolution reconstruction of the cryo-EM density maps.

### Cloning and *in vitro* transcription of *avrE* and *dspE* for oocyte experiments

The *avrE* or *dspE* open reading frame (ORF) was amplified with the following primer sets:

*avrE* forward primer: TTGCCCGGGCGCCACCATGCAGTCACCATCGATCCACCGGA (Kozak sequence underlined),

*avrE* reverse Primer: CCTCTAGATTAGCTCTTCAGTTCGAACCCCTCT

*dspE* forward primer: TTGCCCGGGCGCCACCATGGAATTAAAATCACTGGGAACTG (Kozak sequence underlined),

*dspE* reverse Primer: CCTCTAGATTAGCTCTTCATTTCCAGCCCTTCC

PCR-amplified *avrE* or *dspE* ORF (*Srf*I-*Xba*I fragment) was cloned into *pGH19*^42^ (digested with *Xma*I and *Xba*I) to create *pGH-avrE* or *pGH-dspE*. To prepare cRNA for oocyte injection, *pGH-avrE* or *pGH-dspE* was linearized with *Nhe*I, followed by *in vitro* transcription with T7 polymerase mMESSAGE mMACHINE® Kit (Ambion).

For mutational analysis of DspE, point or deletion mutants of *pGH-dspE* was obtained using Q5 Site-Directed Mutagenesis Kit (New England Biolabs) with following primer sets:

*pGH-dspE*^Δ^*^β-barrel^* (i.e., Δ1278-1566 + Δ1649-1813)

Δ1278-1566 forward primer: GCGGAGCCGGTGACCAGCAACGATA

Δ1278-1566 reverse Primer: ACTGTCCAGGGACAACAGCGTATTC

Δ1649-1813 forward primer: GGAATAGCTCAGGCTAATCCGCAGG

Δ1649-1813 reverse Primer: GCTGTGGCTGTCCGCAGCCTGTTGA

*pGH-dspE^K1399E/K1401E^*

K1399E+K1401E forward primer: CTG<u>G</u>AGTTT<u>G</u>AGCTGACAGAGGATGAG (underline indicate mutation point)

K1399E+K1401E reverse Primer: GCTGTTTTGTAGCGTTCCTTGCAGGGT

pGH-dspE^L1776E/L1777E/L1778E^

L1776E+L1777E+L1778E forward primer: <u>GA</u>G<u>GA</u>A<u>GA</u>GGGGACGAGCAACAGCCTG (underline indicate mutation point)

L1776E+L1777E+L1778E reverse Primer: CGCTGGGGTATTGAAGCCTTCGCTTTT

### *Xenopus laevis* oocyte preparation, injection and expression of DspE and AvrE

Oocytes were purchased as ovary from Xenopus1 Co. (Dexter, MI). The ovary was treated with 0.55 mg/mL of collagenase B (0.191 U/mg) in calcium-free ND96 saline^43^ (96 mM NaCl, 2 mM KCl, 1 mM MgCl_2_, 5 mM HEPES, and 2.5 mM Na-pyruvate, pH at 7.5) for 20 min while on a nutating mixer at 21-22°C. Immediately after treatment, the enzymatic solution was rinsed off from the ovary with ND96 bath saline (96 mM NaCl, 2 mM KCl, 1 mM MgCl_2_, 1.8 mM CaCl_2_, 5 mM HEPES, 2.5 mM Na-pyruvate, and 0.5 mM theophylline, pH at 7.5) several times and cell clusters were spread on several 70 mm plastic culture dishes with ND96 bath saline for temporary storage in an 18°C incubator. On the same day, the follicular cells and follicular membrane covering mature oocytes (stages IV and V) were manually peeled off with fine forceps and oocytes were kept in ND96 bath saline at 18°C until the time for cRNA injection. cRNA was mixed with diethyl pyrocarbonate (DEPC)-treated water to defined concentrations necessary to deliver desired amounts (ranging from 0.01 ng to 20 ng) of cRNA per oocyte when injecting a volume of 27.6 nL. Injection was performed with a nanoinjector (Nanoject II, Drummond Scientific Co. Broomall, PA) following the manufacturer’s directions. DEPC-treated water was injected in control oocytes. Oocytes were kept in a 6-well plastic culture plate at 18°C to allow expression of proteins. Incubation solution was either control (ND96 bath saline), ND96 with inhibitor (PAMAM G0 or G1, niflumic acid, or fipronil), ND96 with 0.0005% fluorescein or ND96 with 0.1% GFP protein.

### Oocyte surface biotinylation assay

Oocytes were injected with 2 ng of wild-type or mutant *dspE* cRNA and incubated in bath ND96 saline for 15 h. Surface-exposed proteins were biotinylated and purified using the Pierce™ Cell Surface Protein Biotinylation and Isolation Kit (ThermoFisher Scientific), following the manufacturer’s protocol with some modifications, as described by Yu et al. 2012^44^. Five oocytes were used per each treatment (with biotin, without biotin or total cell extract). Briefly, cells were rinsed 3 times in OR2 buffer^44^ before incubating in 2.5 mL of OR2 buffer for 10 min with or without Sulfo-NHS-SS-Biotin in 6-well culture plates in a benchtop orbital shaker, set at 85 rpm at room temperature. Oocytes for total cell extract were immediately saved at −80 °C, while oocytes for biotinylation (with or without biotin) were rinsed 3 times in TRIS buffer^44^ before being placed in a 1.5 mL tube with 500 µL of lysis buffer containing 10 µL of Halt™ Protease Inhibitor Cocktail (ThermoFisher Scientific). Lysis mix containing oocytes was homogenized by passing thought a 20-gauge needle for 10 times, before incubation at 4 °C on a nutating mixer. The remaining steps followed the kit manufacturer’s protocol. Final samples were eluted with 200 µL elution buffer and mixed with 50 µL of 5×SDS sample buffer. In parallel, the five oocytes saved for total cell extract were homogenized in 250 µL of 2×SDS sample buffer. For equal loading, 25 µL of total extract or avidin pull down of biotinylated protein samples were added to each lane for SDS-PAGE.

### Two-electrode voltage clamp recordings

Oocytes were injected with 0.01 ng of wild-type or mutant *dspE* cRNA or with 0.1 ng of *avrE* cRNA. AvrE seems less functional in oocytes than DspE. After ∼15h of incubation in bath ND96 saline (96 mM NaCl, 2 mM KCl, 1 mM MgCl_2_, 1.8 mM CaCl, 10 mM HEPES, pH at 7.5), each oocyte was impaled by one voltage-sensing borosilicate microelectrode and one current-passing borosilicate microelectrode with resistance of 0.5±0.1 MOhm, while in 1 mL of an electrically grounded ND96 recording saline (96 mM NaCl, 2 mM KCl, 1 mM MgCl_2_, 1.8 mM CaCl, 10 mM HEPES, pH at 7.5). Note: In initial preliminary experiments, when oocytes were injected with a high amount of *dspE/avrE* cRNA (e.g., 1 ng of *dspE* or 20 ng *avrE* cRNA/oocyte), membrane potentials at 24 h had dropped close to zero mV (Extended Data Table 1) and current conductance was very large (>50 µA), a condition at which the TEVC equipment no longer works properly. Thus, we lowered the cRNA input to 0.01 ng *dspE*/oocyte and 0.1 ng *avrE/*oocyte and evaluation time to 15 h after injection, which yield resting potential similar to that of oocytes injected with water control (Extended Data Table 1) and modest currents that TEVC was able to record.

For ion replacement experiments, variations of ND96 were prepared by replacing the major salt (i.e., 96 mM NaCl) with 96 mM of LiCl, KCl, RbCl, CsCl, Choline-Cl, NDMG-Cl (N-Methyl-D-glucamine Hydrochloride), NaBr, NaI, NaClO_3_, NaBrO_3_ or Na-MES (4-Morpholineethanesulfonic acid, sodium salt)^45,46^. Currents were first recorded in ND96 recording buffer. To replace ND96 with a new cation/anion, 10 mL of a new ND96 solution was slowly added from one end of the recording chamber using a 10 mL plastic syringe with 18G needle, while the original ND96 was washed out using a vacuum outflow 20G tube from the other end of the chamber. The glass microelectrodes were half-filled with 1.5% agar containing 3M KCl. The electrodes were connected to an oocyte clamp amplifier (OC-725C, Warner Instrument Corp. Hamden, CT) by chlorinated silver wires. The bath clamp headstage was connected to bath saline by two chlorinated silver wires inside a disposable polytetrafluoroethylene 18-gauge tubing filled with 1.5% agar containing 3M KCl serving as agar-bridges. The oocyte clamp amplifier was connected to a computer by an analog-digital interface (Digidata 1440A, Molecular Devices, San Jose, CA). The command voltage protocols and data acquisition were performed in Clampex 10.7 (Molecular Devices, San Jose, CA). Oocytes were clamped to a desired potential using the fast clamp mode with maximum clamp gain and current gain set to 0.1 V/uA. Signal for both voltage and currents was recorded. Upon impaling an oocyte and before clamping it, both electrodes are capable of measuring the resting potential of that oocyte. Oocytes were clamped at their resting potential and test pulses of 100 ms toward more positive or more negative potentials in 10 mV increments were applied. The resultant current was recorded and analyzed. Current amplitude was determined at 10 to 20 ms of the test pulse, time at which there was the smallest or no overlaps with membrane capacitance or Ca-dependent chloride currents from endogenous oocyte channels^47^. Since resting potential values across individual oocytes varied (likely due to uncontrollable intrinsic differences in each oocyte, its size and in fine adjustment of electrode position and resistant), the voltage-current relationship data was fit to a quadratic polynomial regression (SigmaPlot 12.5 Systat Software Inc.) providing intermediate values and 95% confidence intervals. This also allowed currents elicited by the test potentials on control oocytes to be subtracted from the currents in treatment oocytes, so the resultant values represent only DspE- or AvrE-mediated current flowing across the membrane. Comparison of the currents was performed using a two-way ANOVA with Tukey’s test, with significance set to a *P* value < 0.05.

### Oocyte swelling assay

Oocytes injected with 1 or 2 ng *dspE* or 20 ng *avrE* cRNA/oocyte were imaged using Motic Images Plus 3.0 software connected to a Moticam X3 camera (Motic China Group Inc., China) on a SHR Plan Apo 1X WD:60 magnification lens of a stereoscope (Nikon SMZ18 Nikon Corp., Japan). At 0.01 ng *dspE* or 0.1 ng *avrE* cRNA/oocyte that was used for TEVC recordings, no baseline oocyte swelling was observed. Baseline swelling began to be observed at >0.1 ng *dspE* or >10 ng *avrE* cRNA/oocyte. For baseline oocyte swelling, starting oocyte images were recorded immediately after each injection, and then every 2 h to 4 h intervals for 24 h. Oocytes were kept in bath saline of 200 mOsm with or without PAMAM inhibitors in the stereoscope room at 18-19°C for the entire period. Each picture depicting 5 oocytes (replicates) was analyzed with the Fiji software^48^. Data are presented as absolute volume at a given evaluation time or as change in volume in relation to the start point (immediately after cRNA injection). For hypoosmotic-induced swelling, oocytes expressing DspE or AvrE were transferred into a 5-fold diluted ND96 bath saline (40 mOsm) and were immediately imaged as described for baseline swelling once every 20 s for 10-20 min or until oocytes injected with *dspE* or *avrE* started to burst. Data are presented as change in volume in relation to the first picture in diluted saline. Pictures were also arranged in sequence to create time-lapse movies showing oocyte swelling and bursting. One-way ANOVA with Tukey’s test was used for multiple comparisons within a data set, with significance set to a *P* value < 0.05. For dataset with repeated measures over time, as in the hypoosmotic-induced swelling assay, a two-way repeated measure ANOVA with Dunnett’s test was used instead, with significance also set to a *P* value < 0.05.

### Oocyte dye uptake assay

Two hours after injection with 1 or 2 ng of *dspE* or 20 ng of *avrE* cRNA, oocytes were placed in ND96 bath saline with or without 5 µg/mL fluorescein, 1 mg/mL GFP and/or 5 mM PAMAM G1 inhibitor and incubated until evaluation time, as indicated in figure legends. Oocytes were rinsed twice in ND96 bath saline and imaged as described above for oocyte swelling assay, with a few exceptions: It was imaged at a 2× magnification with either a bright field or GFP-B filter. In the Motic Images Plus software, the green channel gain was increased to improve green fluorescence detection. While specific values of the green channel gain value varied across different independent assays, all configurations were kept the same across all treatments within the same experiment. Bright field and fluorescent images of each oocyte were stacked using the Fiji software and the integrated density of fluorescence was measured within oocyte boundaries and subtracted from the background, so data is presented as corrected total cell fluorescence (for short: cell fluorescence). Two-way ANOVA, with Tukey’s test, was used for multiple comparisons within a dataset, with significance set to a *P* value < 0.05.

### eGFP purification

For eGFP purification, *pET28-eGFP*^49^ was transformed into *E*. *coli* Rosetta(DE3). eGFP production was induced by adding 0.25 mM IPTG to LB bacterial culture for 4 h at 28 L. eGFP was purified from total cell lysate using Ni-NTA agarose beads in the extraction buffer (50 mM Tris-Cl, pH 8.0, 250 mM NaCl, 5 % Glycerol, 0.1 mM PMSF). Before oocyte uptake test, buffer was exchanged to ND96 bath saline using Amicon Ultra-4 Centrifugal Filter Units (Millipore Sigma).

### Western blot analysis

Five oocytes (15 mg) or 10 mg fresh plant leaf tissue was homogenized in 100 µL of 2×SDS sample buffer. After 10 min boiling, cell lysates were brief centrifuged and 10 µL was loaded to each lane of an SDS-PAGE gel. After separation, proteins were blotted onto a PVDF membrane. AvrE, β-Actin, DspE, or PR1 was detected by anti-AvrE^20^, anti-beta Actin [HRP] (GenScript), anti-DspE^50^, anti-PR1 antibody (a gift from Xinnian Dong), respectively.

### Liposome preparation and liposome dye release assay

Soy Extract Lipids in chloroform were purchased from Avanti Polar Lipids and stored in glass vials^24^. The lipids dissolved in chloroform was evaporated under a stream of nitrogen until it forms a thin lipid film and then dried in a vacuum desiccator chamber overnight. On the second day, the lipid film was dissolved in a suspension buffer (HBS buffer: 20 mM HEPES, 300 mM NaCl, pH 8.0) containing 50 mM 5(6)-carboxy-fluorescein (CF; Novabiochem^®^) or 50 mg/mL Polysucrose 40-fluorescein isothiocyanate conjugate (FITC-polysucrose, MW of 30-50 kDa with an estimated diameter of 80 Å, Millipore Sigma). To solubilize lipids, the solution in the glass vial was sonicated for 15 min first and then incubated in a 37 °C water bath for at least an hour. Then the lipid solution was subjected to freeze-thaw cycles for 8 times, in which lipids were frozen in liquid nitrogen for 5 min and then thawed in 37 °C water bath for 10 min, to reduce the formation of multilamellar liposomes. To control the liposome size, liposomes were extruded through a polycarbonate filter (200 nm, Whatman) 25 times using a mini extruder (Avanti Polar Lipids) with Hamilton glass syringes. CF- or FITC-polysucrose-loaded liposomes were purified by centrifugation at 41,000 rpm for 20 min in a TLA 100.3 rotor incorporating three sequential wash steps. After the final wash, CF- or FITC-polysucrose-loaded liposomes were resuspended in HBS buffer to make the final CF-loaded liposome concentration of 1 mg/mL and FITC-polysucrose-loaded liposome concentration of 0.5 mg/mL^51^.

Release of the liposome contents was assessed using the self-quenching property and fluorescence of CF and FITC-polysucrose. The HBS buffer composition in and outside the liposome is the same (20mM HEPES, 300mM NaCl, pH8.0). Permeability induced by DspE was evaluated by incubating 10 µL DspE protein solution with 90 µL carboxyfluorescein- or FITC-polysucrose-loaded liposomes (0.25 μg/μL). The fluorescence intensity was measured every 30 s continuously for 2 hours after addition of the purified DspE protein (WT or mutant) to the liposomes. Then 5 µL of 20% Triton X-100 (Sigma-Aldrich) was added into the 100 µL solution to fully release the dye and its readings were measured for 20 min. The average reading of the last three minutes was used for normalization (100% dye release). In the compound inhibition assays, the buffer, DspE protein, and PAMAM G1 inhibitor, at a total volume of 10 µL were first mixed thoroughly with pipetting, then 90 µL CF- or FITC-polysucrose-loaded liposomes were added to make the total volume 100 μl. The spectrofluorometric excitation and emission parameters were set at the wavelengths of 485 and 510 nm for CF/FITC-polysucrose molecules.

The DspE protein stock solutions (2.5-25 μM) contained 1 mM DTT. The liposome assays were done at 0.05-0.15 μM DspE concentrations with the final DTT concentration less than 0.02 mM. The presence of DTT in the protein buffer did not affect the fluorescence of carboxyfluorescein (Extended Data Fig. 5e). Similarly, the presence of PAMAN G1 at concentrations from 0.3 – 300 μM did not affect the intrinsic fluorescence of carboxyfluorescein (Extended Data Fig. 7e).

### Bacterial media and plant growth

Bacterial strains used are wild-type *Pseudomonas syringae* pv. *tomato* strain DC3000 and its mutants: the ΔE mutant^11^, ΔM mutant^11^ and the ΔEM mutant^11^ and wild-type *Erwinia amylovora* strain Ea273 and its *dspE* mutant^8^. Bacteria were grown in low-salt Luria-Bertani (LB) medium at 28°C. Antibiotic ampicillin, gentamicin, kanamycin, rifampicin or spectinomycin was added at 200, 10, 50, 100 or 50 µg/ml, respectively. *Arabidopsis thaliana* Col-0 and Col-0/*DEX::his-avrE*^20^ plants were grown in Redi-Earth potting soil (Sun Gro Horticulture) in air-circulating growth chambers. Plants were grown under relative humidity set at 60%, temperature at 20°C, light intensity at 100 µE m^2^ s and photoperiod at an 8 h light-16 h dark cycle. Four- to five-week-old plants were used for bacterial disease assay. Immature pear fruits were gifts from George Sundin at Michigan State University. *Nicotiana benthamiana* plants were grown in a growth chamber with 12 hr light, 12 hr dark at 23°C day/21°C night, ∼55% humidity and ∼100 μmol/m^2^/s light intensity. Four- to six-week-old plants were used for transient expression assay.

### Bacterial disease assays

Disease assays with immature pear fruits were performed as previously reported^8^. Pears were surface-sterilized with 10% bleach for 5 min and rinsed in sterile water twice. Then, a small hole was made in the pear using a 200 µL tip. Ten µL of 10^3^ CFU/mL Ea273 or the *dspE* mutant was loaded into the hole. Inoculated pears were placed on a wet paper towel in a sterile box to maintain high humidity at 28L for 10 days. Disease assays with Arabidopsis plants were performed as follows. Arabidopsis plant leaves were infiltrated with *Pst* DC3000, ΔE, ΔM or ΔEM at 10^6^ CFU/ml with a needle-less syringe. After visible water soaking disappeared (within 1 h), plants were kept under high humidity (∼99%) at 23°C. Bacteria population in leaves was determined at day 3 post infiltration. Detached leaves were surface-sterilized in 75% ethanol for 30 s and rinsed in sterile water twice. Then, leaf discs (1 cm^2^ in diameter) were punched out and ground in 100 µL sterile water. Ten µL of each 10-fold serial diluted leaf extract was plated on LB rifampicin and kept at 28°C for 24 h. Colony forming units (CFUs) were counted under microscope before colonies started to coalesce and analyzed by GraphPad Prism software. Two-way ANOVA with Tukey’s test was used for multiple comparisons within a data set, with significance set to a *P* value < 0.05. For inhibition assays, 50 nM PAMAM G1 was added to bacterial suspension and co-inoculated into plants.

### AvrE/DspE sequence alignments

Sequences of *E. amylovora* DspE, *P. carotovorum* DspE, *Pst* DC3000 AvrE and *P. stewartia* WtsE are aligned using Clustal Omega^52^. Sequences are entries from uniprot (https://www.uniprot.org) as listed below:

*Erwinia amylovora* Ea321 DspE: https://www.uniprot.org/uniprotkb/O54581/entry

*Pectobacterium carotovorum* Er18 DspE: https://www.uniprot.org/uniprotkb/D5GSK5/entry

*Pseudomonas syringae* pv. tomato DC3000 AvrE: https://www.uniprot.org/uniprotkb/Q887C9/entry

*Pantoea stewartii* subsp. *stewartii* SS104 WtsE: https://www.uniprot.org/uniprotkb/Q9FCY7/entry

### Transient expression of DspE in *Nicotiana benthamiana*

*dspE* and *dspE* mutant ORFs were PCR-amplified with *dspE* ORF forward primer (TT<u>GGGCCC</u>ATGGAATTAAAATCACTGGGAACTG, underline indicates *Apa*I site) and *dspE* ORF reverse primer (TTT<u>ACTAGT</u>TTAGCTCTTCATTTCCAGCCCTTCC, underline indicates *Spe*I site) and *pGH-dspE* or *pGH-dspE* mutant plasmids as template. PCR-amplified *dspE* and *dspE* mutants ORF (*Apa*I-*Spe*I fragment) were cloned into the binary vector *pER8*^53^ to create *pER-dspE* and *pER-dspE* mutant constructs. All constructs were transformed into *Agrobacterium tumefaciense* GV3101 for transient expression assay. 1×10^8^ CFU/mL of *A. tumefasciens* GV3101 containing *pER8* empty vector, *pER-dspE* or *pER-dspE* mutant were syringe-inoculated into leaves of *N. benthamiana* and kept at 22L for 24 h before leaves were painted with 90 µM estradiol. Eight hours later, leaf samples were collected for western blot. Water-soaking/necrosis symptoms were recorded at 8 h and 24 h after estradiol treatment under high humidity (>95%).

### Arabidopsis leaf protoplast swelling assay

Leaf mesophyll protoplasts were isolated from 5 week-old Arabidopsis Col-0 and transgenic Col-0/*DEX::his-avrE*^20^ following the Tape Sandwich method^54^. For swelling test, isolated protoplasts were incubated in protoplast isolation medium (MMg), which contains 400 mM mannitol (isosmotic), or in MMg containing 320 mM mannitol (hypoosmotic) for 1 h. Protoplast images were taken using Leica DM500 microscope with ICC50W camera. Protoplast volumes were analyzed with Image J software.

### Callose Staining

Callose staining was performed as described previously^13^. Callose images were taken using a Zeiss Axiophot D-7082 Photomicroscope. The number of callose depositions was determined with Quantity One Colony Counting software (Bio-Rad).

### Statistical analysis

Experimental sample size was chosen based on previously published literature to be sufficient for statistical analyses. Three to four plants (biological replicates) per treatment and/or per genotype were analysed per individual experiment. Two or more independent experiments were performed for all assays. The following statistical analyses were employed: (1) one-way analysis of variance (ANOVA) was used for multi-sample experiments with one variable, followed by Tukey’s honest significant difference (HSD) test for multi-comparisons; (2) two-way ANOVA was employed for multi-variable analyses, followed either by Tukey’s HSD test for multi-comparisons or by Dunnett’s test for comparison against a common control treatment; (3) two-way repeated measure ANOVA was used for repeated measures over same experimental unit, followed by Dunnett’s test for comparison against a common control treatment; and (4) Student’s t-test was used to compare two sets of data. If normality of the residues and equality of variances test failed, non-parametric alternatives ANOVA on Ranks or Mann-Whitney Rank Sum Test was used instead. All statistical tests are described in the figure legends, methodology and source data files. Graphic plots were generated by SigmaPlot 12.5 and show mean ± SEM and individual data points.

### Graphic design

Images and cartoons were created or assembled in CorelDRAW v.22 (Corel Corp. Ottawa, Canada). All graphics were first plotted on SigmaPlot 12.5 (Systat Software Inc. San Jose, CA) and then further edited for color and arrangement in the figure panels in CorelDRAW v.22 (Corel Corp. Ottawa, Canada).

## DATA AVAILABILITY

Data needed to evaluate this paper are available in the main text and Supplementary Information. Uncropped gel and blot source data are provided in Supplementary Figures 1–3. Source data (with statistical analyses) for Figures 1–4 and Extended Data Figs. 1– 10 are provided with this paper. Gene and protein sequence data were obtained from uniprot (https://www.uniprot.org).

## CODE AVAILABILITY

No customized code was generated in this study. All bioinformatic tools and software used in this study are cited in the text.

### ACKNOWLEDGEMENTS

We sincerely thank the following colleagues for sharing biological materials: Immature pear fruits were gifts from Dr. George Sundin at Michigan State University. The ΔE mutant, ΔM mutant and the ΔEM mutant of *Pseudomonas syringae* pv. tomato DC3000 were from Dr. Alan Collmer at Cornell University. *Erwinia amylovora* strain 273 and its *dspE* mutant from the late Prof. Steven Beer at Cornell University. We thank Duke Phytotron for technical assistance. This work was supported by NIAID 1R01AI155441 (S.Y.H.), Duke Science and Technology Initiative (K.D. and S.Y.H.), and NIGMS GM145026 (P.Z.).S.Y.H. is an investigator at the Howard Hughes Medical Institute.

## AUTHOR CONTRIBUTIONS

K.N., P.Z. and S.Y.H. conceived the project and designed the initial experiments. K.N., F.A., J.C. and P.Z. performed most of the experiments. F.A. and K.N. were involved in oocyte swelling and fluorescence dye uptake assays. P.Z. and J.C were involved in model building and liposome permeability assay. F.A. was involved in oocyte TEVC assays. K.D., P.Z. and S.Y.H. supervised the project team. K.N., F.A., J.C., K.D., P.Z. and S.Y.H. wrote the manuscript.

## COMPETING INTERESTS

The authors declare no competing interests.

## ADDITIONAL INFORMATION

Correspondence and requests for materials should be addressed to Sheng Yang He, Pei Zhou and Ke Dong. Reprints and permissions information is available at www.nature.com/reprints.

## EXTENDED DATA AND LEGENDS

**Extended Data Table 1.** Effects of DspE/AvrE expression on oocyte membrane resting potential.

| Evaluation time<br>(h after injection) | Treatment | Oocyte resting potential (mV) |  |
| --- | --- | --- | --- |
|  |  | mean (n=5) | SEM |
| 24 | H<sub>2</sub>O | -22.04 | 0.776 |
|  | DspE 2 ng/cell | -0.18 | 0.442 |
| 15 | H<sub>2</sub>O | -23.46 | 1.655 |
|  | DspE 0.01 ng/cell | -23.22 | 0.198 |
| 24 | H<sub>2</sub>O | -23.24 | 1.013 |
|  | AvrE 20 ng/cell | -0.1 | 0.325 |
| 15 | H<sub>2</sub>O | -16.72 | 0.527 |
|  | AvrE 0.1 ng/cell | -16.6 | 0.394 |

Experiments were independently performed three times with similar results. Exact P-values for all comparisons are detailed in the Source Data files.

**Extended Data Figure 1.**
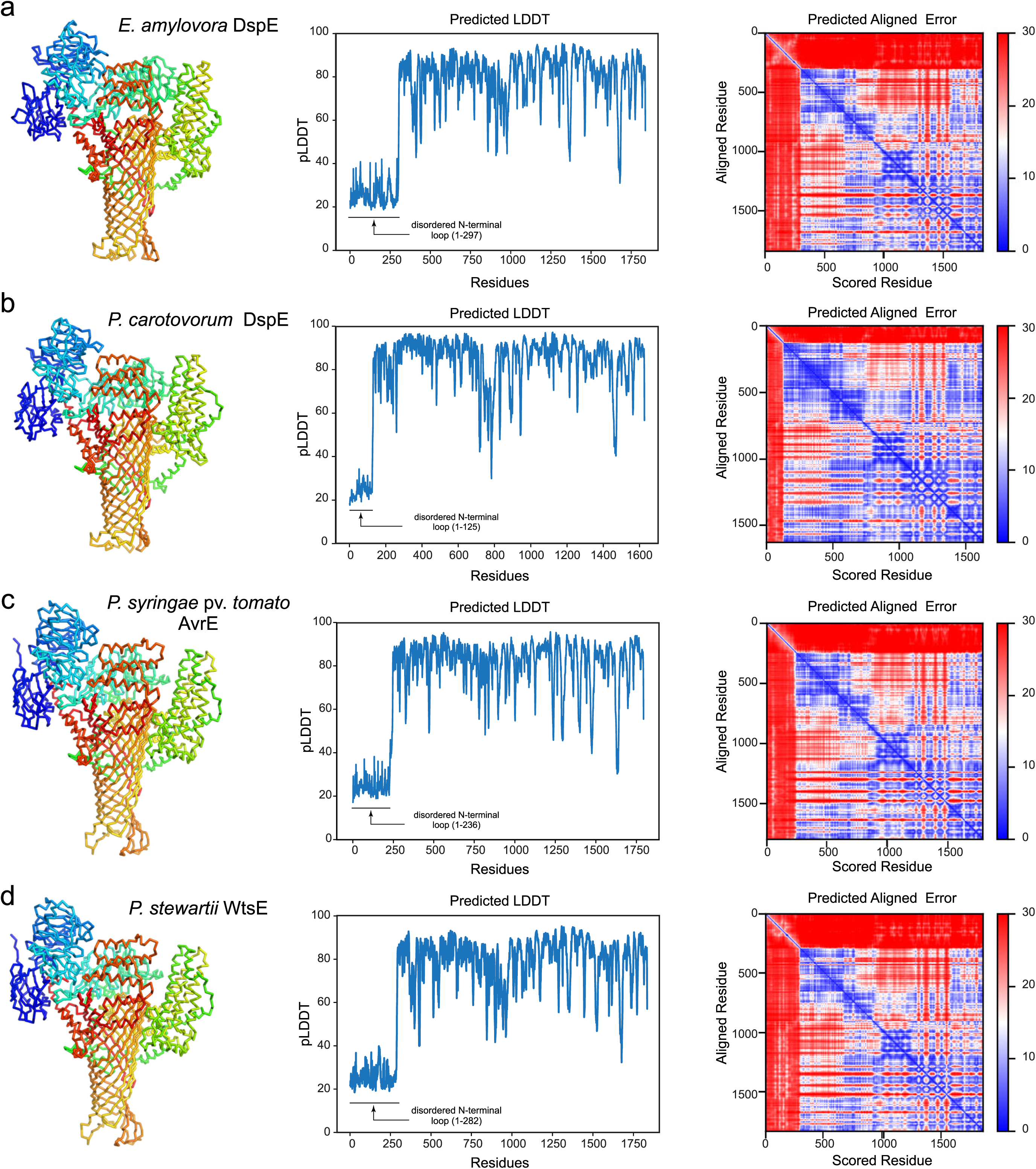
AlphaFold2 models of the AvrE/DspE-family effector proteins. The top-ranked models, predicted local distance difference test (pLDDT), and predicted aligned error (PAE) of *E. amylovora* DspE, *P. carotovorum* DspE, *P. syringae* pv. tomato AvrE, and *P. stewartii* WtsE are shown in panels **a**, **b**, **c**, and **d,** respectively. The models are predicted by AlphaFold2 using MMseqs2 (ColabFold). For each protein, the N-terminal disordered loop with low pLDDT scores is not shown, while the remaining residues are shown in Ca traces and colored in rainbow, with the N-terminus in blue and C-terminus in red.

**Extended Data Figure 2.**
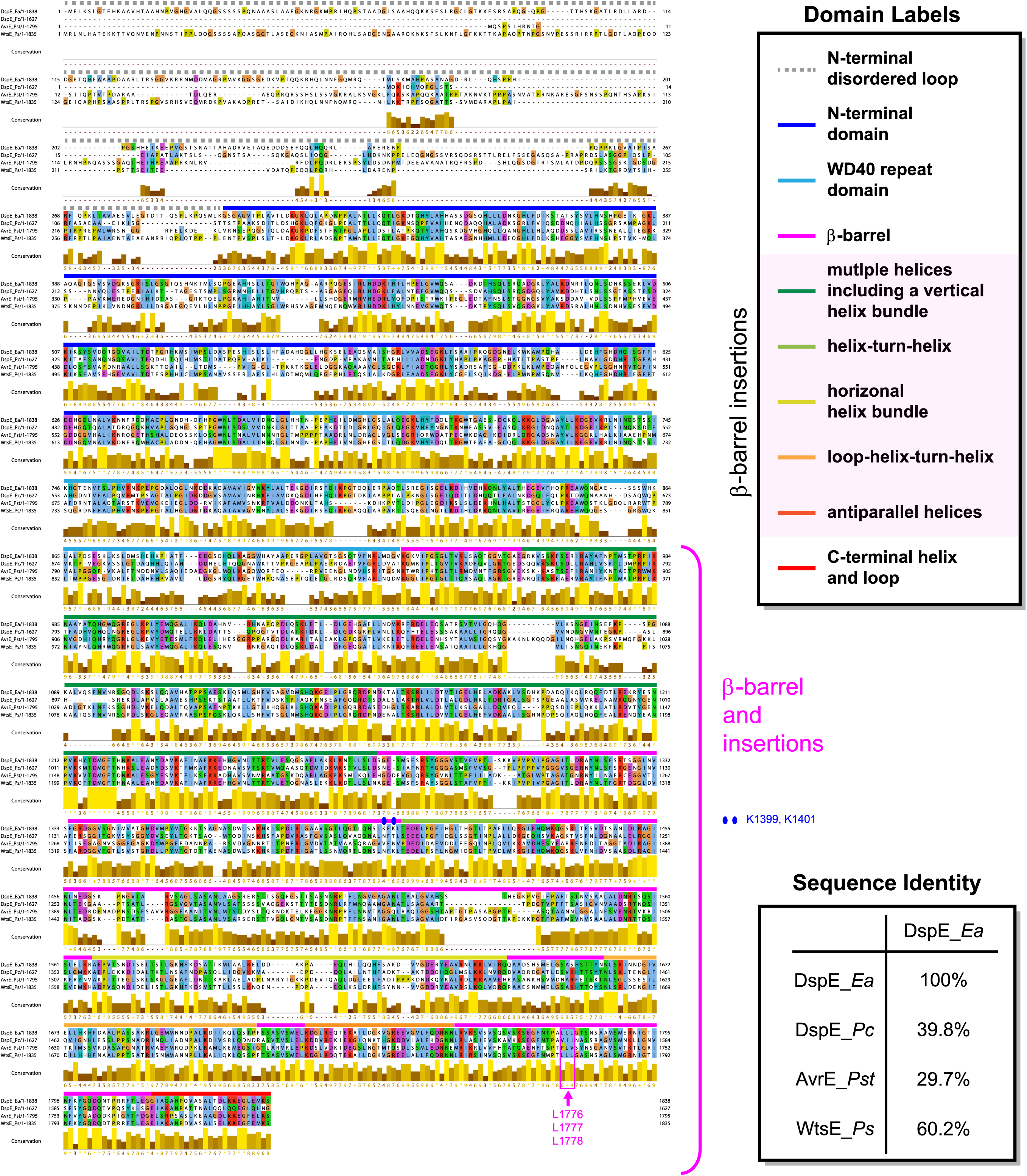
Sequence alignment of representative AvrE/DspE-family effectors. Sequences of *E. amylovora* DspE, *P. carotovorum* DspE, *Pst* DC3000 AvrE, and *P. stewartii* WtsE are aligned using Clustal Omega^44^. Domain regions are indicated above the aligned sequences. Sequence identities between different protein pairs are labeled. The β-barrel is formed with multiple helices and long loops inserted within the primary sequence of the β-barrel. In *E. amylovora* DspE, the β-barrel starts from a small β-hairpin from K932-E956, which is followed by a large insertion of multiple helices (including a vertical helix bundle) from E957-D1276. The main β-barrel follows from S1277-G1813, though it is disrupted by several insertion loops and helices, including T1403-E1430 (a helix-turn-helix motif), A1567-H1647 (a horizontal helix bundle), N1662-P1712 (an insertion loop followed by a helix-turn-helix motif), and K1723-N1753 (two antiparallel helices). The β-barrel is further appended with a C-terminal helix and an extended loop (G1814-S1838). Mutated outward-facing residues (L1776, L1777, and L1778) and inward-facing residues (K1399 and K1401) of the β-barrel are labeled in pink and blue, respectively.

**Extended Data Figure 3.**
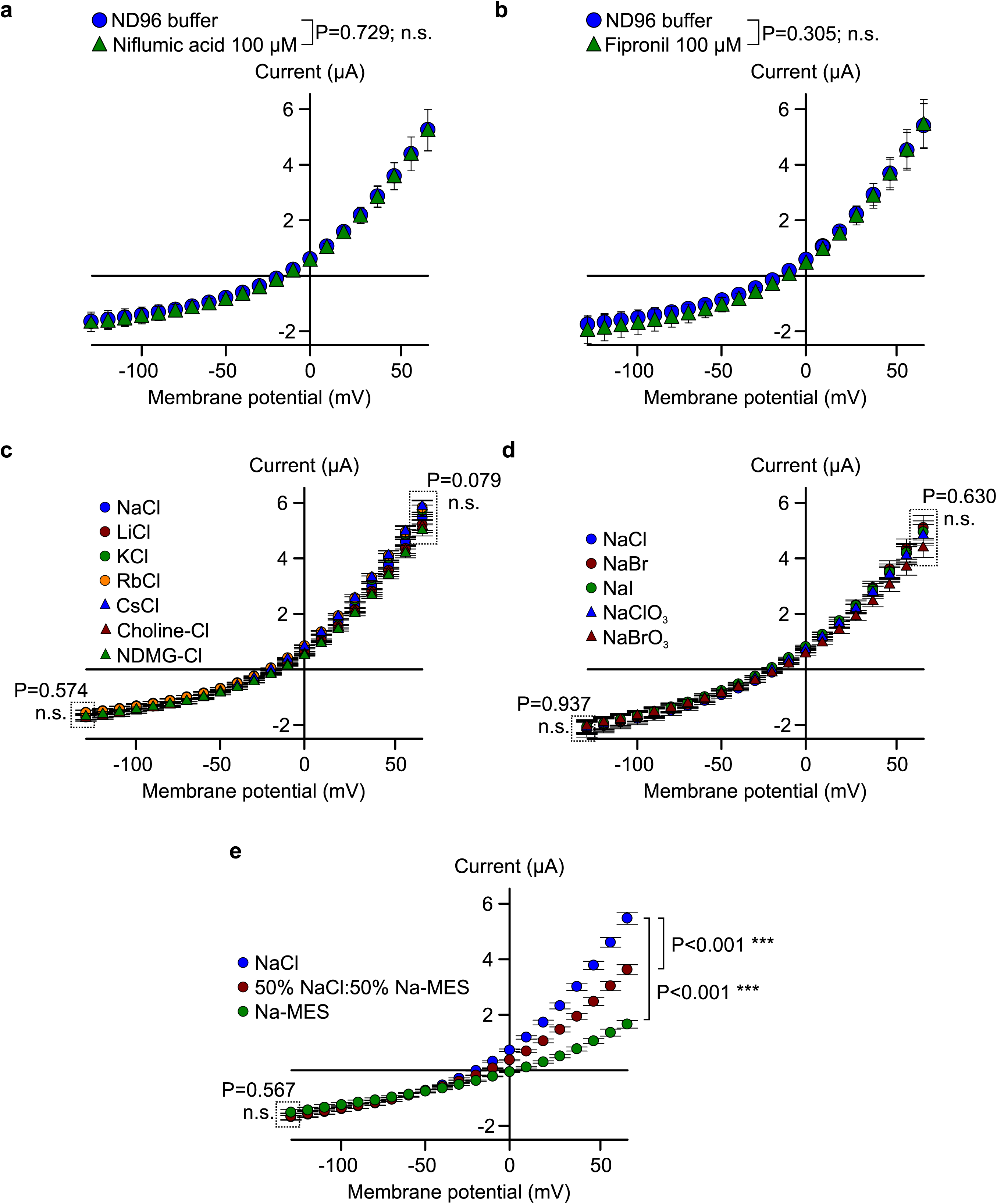
Characterization of DspE induced whole-cell currents. **a**,**b**, DspE-induced currents were not inhibited by niflumic acid or fipronil. Current values (mean ± SEM; n=5) at different test pulses from oocytes expressing DspE proteins were recorded once in ND96 recording buffer at 15 h after injection with 0.01 ng cRNA, and a second time after 10 min of incubation with 100 µM of each inhibitor. **c**, Cation replacement experiment. After 15 h of incubation, cells were recorded in normal ND96 buffer, and then in a new recording solution where sodium in ND96 was replaced with another cation (see details in the Methods section). The data show mean ± SEM (n=5) values at each test pulse for each cation after subtracting currents from control cells. **d**,**e**, Anion replacement experiment. Same as presented in c, but with chloride in ND96 being replaced by other elementary anions (d) the organic anion MES^−^ (e). In **e**, either 100% or only 50% of the Cl^−^ was replaced by MES^−^. Experiments were independently performed three (a,c,e) or two (b,d) times with similar results. Two-way ANOVA values and exact P-values for all comparisons are detailed in the Source Data files.

**Extended Data Figure 4.**
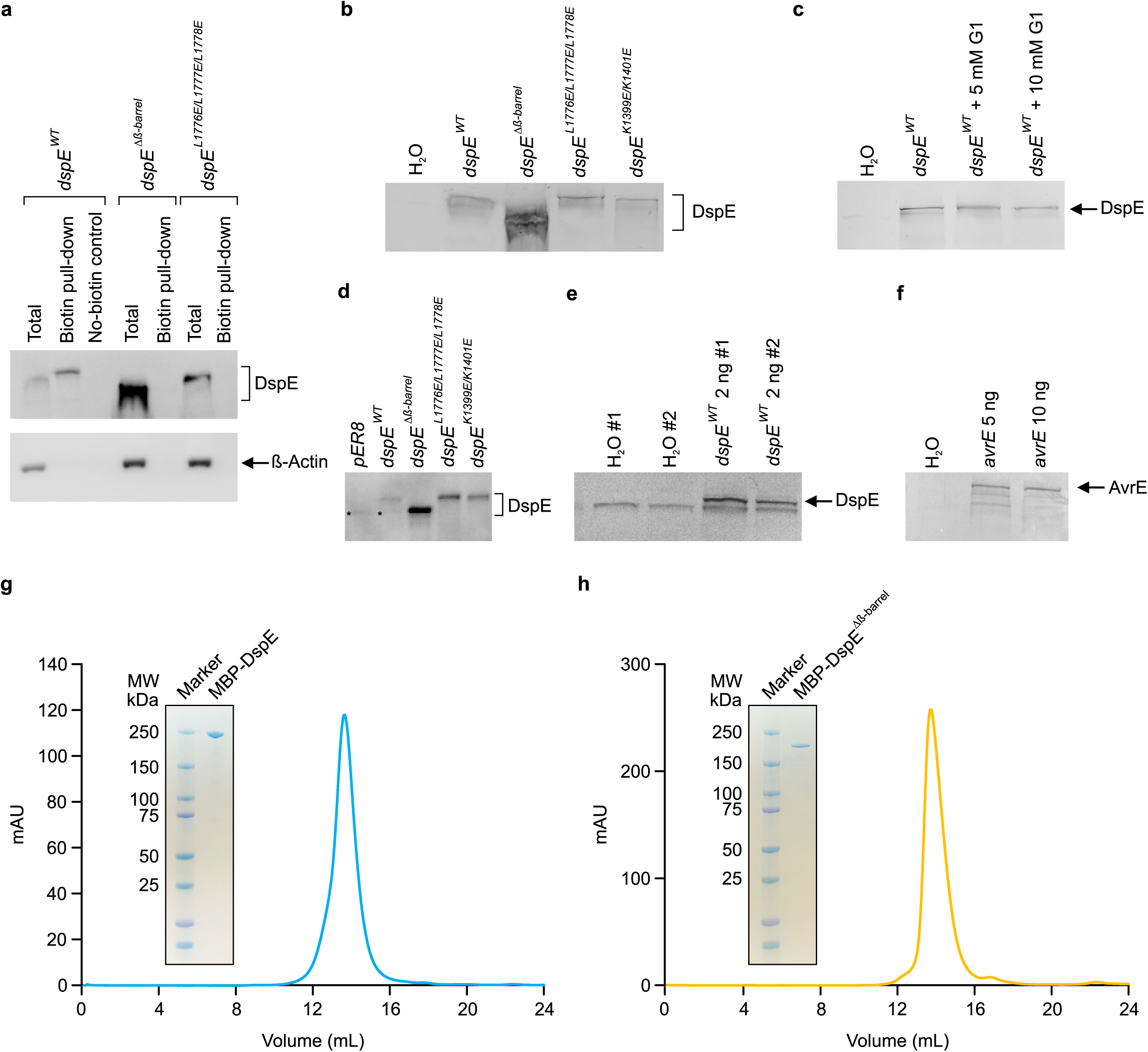
Immunoblot detection of AvrE and DspE proteins expressed in oocytes and in tobacco leaves and elution profile of MBP-DspE and MBP-DspE^Δβ-barrel^ proteins. **a**, Oocyte surface biotinylation assay of DspE. Two nanograms of wild-type or mutant *dspE* cRNA was injected into oocytes, which were incubated in ND96 media at 18°C for 15 h before being subjected to surface protein biotinylation assay (see Methods). Anti-β-Actin antibody detect β-Actin, which is a cytoplasmic protein, serving as a negative control. **b**,**c**,**e**,**f**, Detection of AvrE, DspE and/or mutant DspE proteins expressed in oocytes. Oocytes injected with 1 ng (unless otherwise noted in the gel lane label) *avrE*, *dspE* or mutant *dspE* cRNA or H_2_O (control) were incubated in ND96 media at 18°C for 19 h before being subjected to SDS-PAGE and immunoblot analysis. In **c**, oocytes were incubated in absence or in presence of 5 or 10 mM of the inhibitor PAMAM G1. **d**, Detection of DspE proteins expressed in tobacco leaves. 1×10^8^ CFU/mL of *Agrobacterium tumefaciens* GV3101 containing *pER8* empty vector, *pER8-dspE* or *pER8-dspE* mutants were syringe-inoculated into leaves of *Nicotiana benthamiana* and kept at 22°C for 24 h before leaves were painted with 90 μM estradiol to induce protein expression for 8 h before subjected to SDS-PAGE and immunoblotting. Asterisks in the *pER8* and DspE^WT^ lanes indicate a faint nonspecific protein band. Because active DspE expressed much more poorly in plant cells than mutant DspE proteins, mutant DspE samples were diluted 20 times with 2×SDS sample buffer (See Supplementary Figure 1d). Experiments were independently performed two times. See Supplementary Figure 1 for image cropping. **g,h**, Purified WT MBP-DspE (g) or MBP-DspE^Δβ-barrel^ (h) was eluted on a Suprose 6 increase 10/300 GL column and was analyzed on a SDS-PAGE gel. FPLC traces and gel images in g and h are representative of 3 experimental replicates. See Supplementary Figure 3 for image cropping.

**Extended Data Figure 5.**
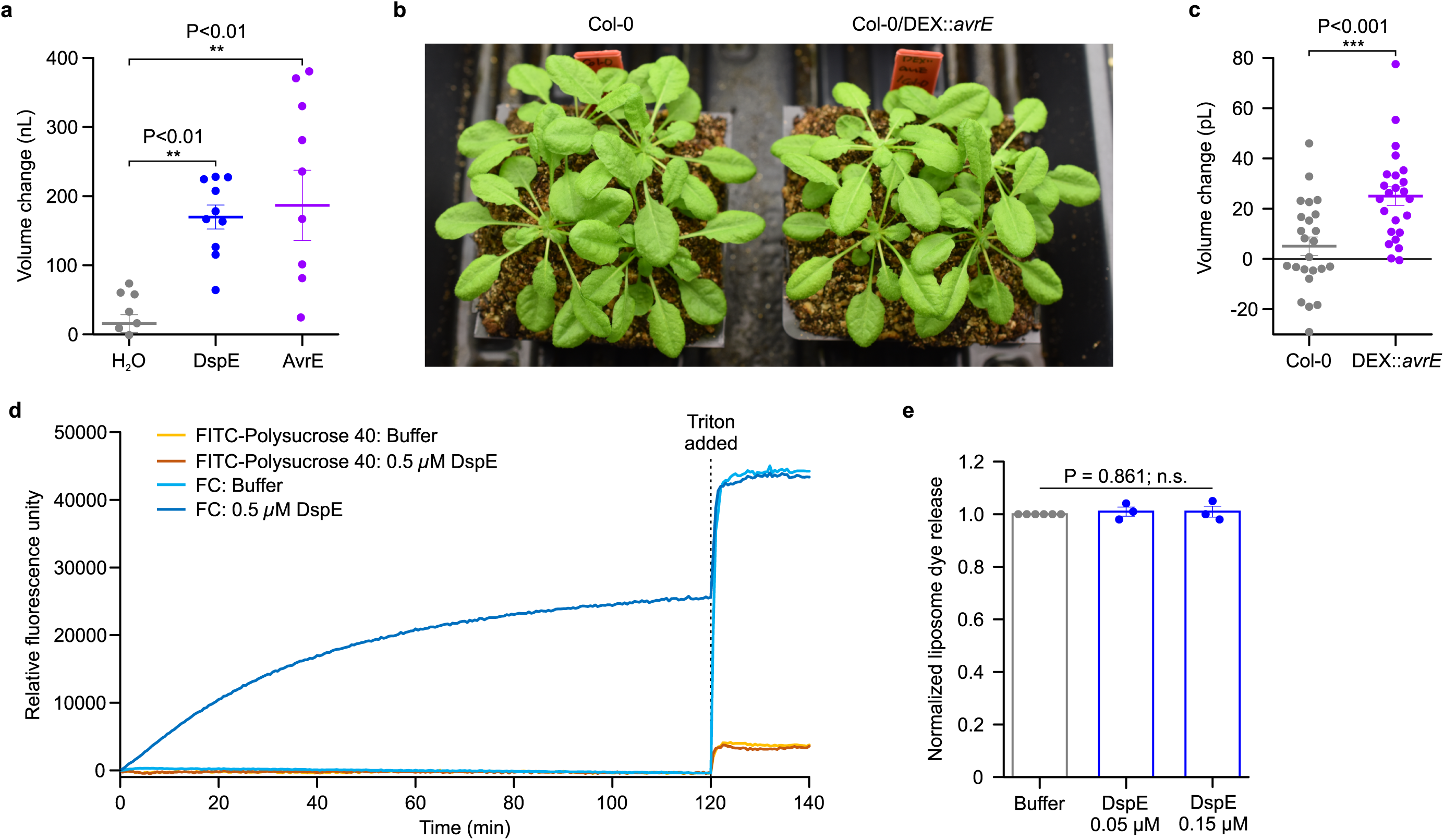
DspE activities in *Xenopus* oocytes, Arabidopsis and liposome. **a,** DspE and AvrE induced a baseline swelling of many oocytes in 200 mOsm bath saline. Oocytes were imaged and measured at 24 h after 2 ng (*dspE*) or 20 ng (*avrE*) cRNA injection. Plot shows mean ± SEM (n=10) and individual replicate values for the increased oocyte volume in relation to its initial volume. See Extended Data Figure 4e,f for immunoblotting of DspE and AvrE proteins expressing in oocytes. **b**, Five-week-old Arabidopsis wild type Col-0 and transgenic Col-0/*DEX::his-avrE* plants (basal expression without dexamethazone induction). **c,** Changes in protoplast volume (mean ± SEM; n=24) when isolated protoplasts were incubated in protoplast incubation buffer containing 320 mM (low osmolarity) mannitol for 1 h compared to protoplast incubation buffer containing normal 400 mM mannitol for 1 h before image capture and volume analysis with Image J software. **d**, Liposome dye release assay using fluorescein isothiocyanate conjugated polysucrose 40 (FITC-polysucrose 40) and carboxyfluorescein (CF). **e**, Normalized liposome dye release (mean ± SEM; n=3) after addition of triton. Raw fluorescence readings were normalized to the buffer control samples. Experiments presented in this figure were independently performed two times. One-way ANOVA on Ranks (a,e) or two-tailed student’s *t*-test (c) values and exact P-values for all comparisons are detailed in the Source Data files.

**Extended Data Figure 6.**
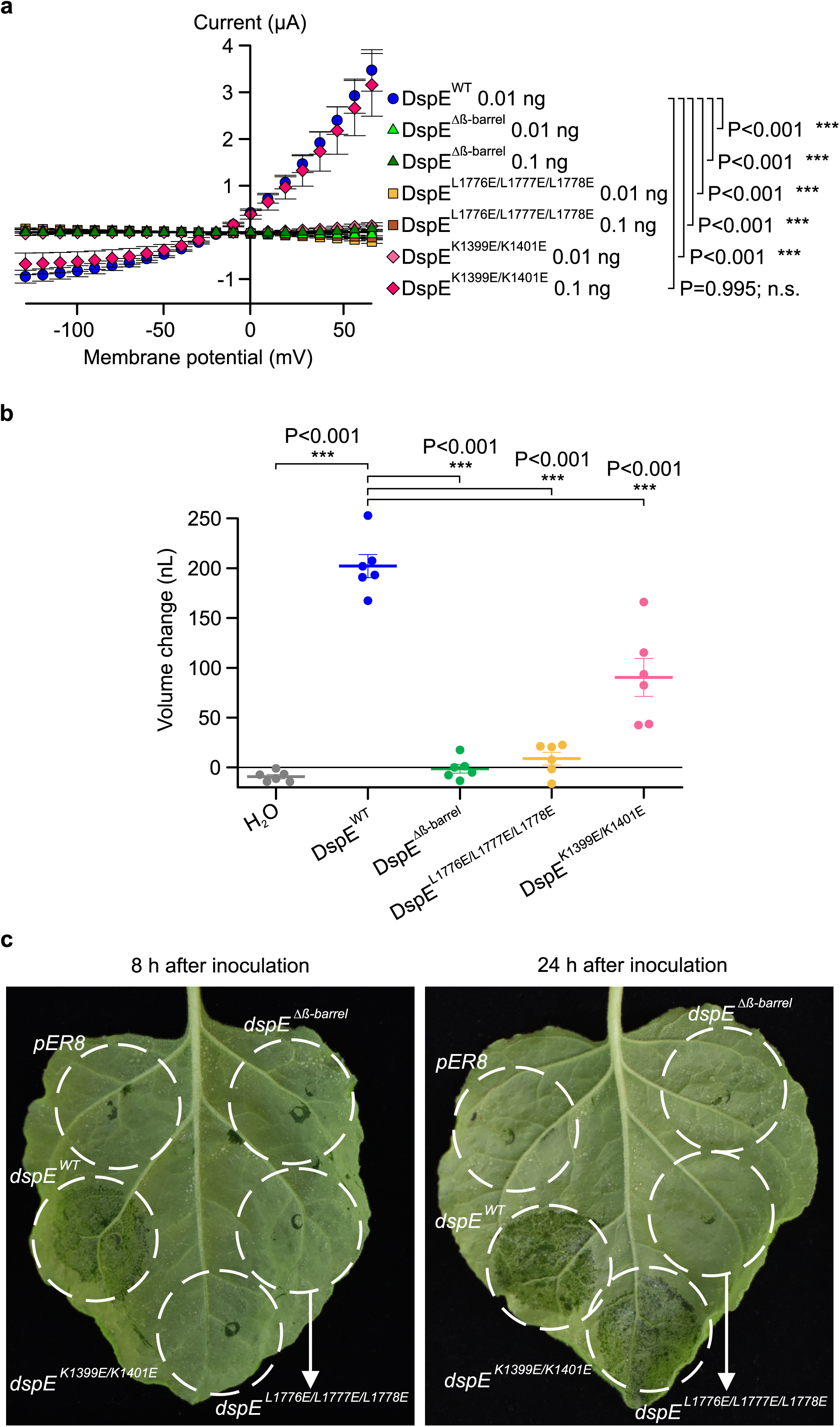
Functional analysis of mutant DspE proteins in Xenopus oocytes and tobacco. **a**, Effect of mutations on DspE-induced currents in oocytes. Mean ± SEM (n=5) current values at different test pulses from oocytes expressing wild-type or mutant DspE proteins at 15 h after injection with 0.01 or 0.1 ng cRNA. Note: At 0.01 cRNA injection, all mutant DspE proteins did not induce currents. Next, 10-fold increase in mutant *dspE* cRNA (i.e., 0.1 ng) was injected into oocytes, revealing current induction by DspE^K1399E/K1401E^, suggesting the K1399E/K1401E double mutations only partially affect ion conductance, consistent with results in panels b and c. **b**, Effect of mutations on DspE-induced baseline swelling in oocytes. Plots represent mean ± SEM (n=6) and individual values of increased oocyte volume in relation to its initial volume 15 h after 1 ng cRNA injection. **c**, Water-soaking assay in tobacco leaves. 1×10^8^ CFU/mL of *Agrobacterium tumefaciens* GV3101 containing *pER8* empty vector, *pER-dspE*, *pER-dspE* mutants were syringe-inoculated into leaves of *Nicotiana benthamiana* leaves (infiltration areas circled) and kept at 22L for 24 h before leaves were painted with 90 µM estradiol to induce DspE expression. Water-soaking symptom (dark-coloured appearance) was assessed at 8 h and 24 h after estradiol induction, showing a complete loss of water-soaking induction by DspE^β-barrel^ and DspE^L1776E/L1777E/L1778E^, but only delayed water-soaking by DspE^K1399E/K1401E^. Experiments were independently performed two times. Two- (a) or one- (b) way ANOVA values and exact P-values for all comparisons are detailed in the Source Data files.

**Extended Data Figure 7.**
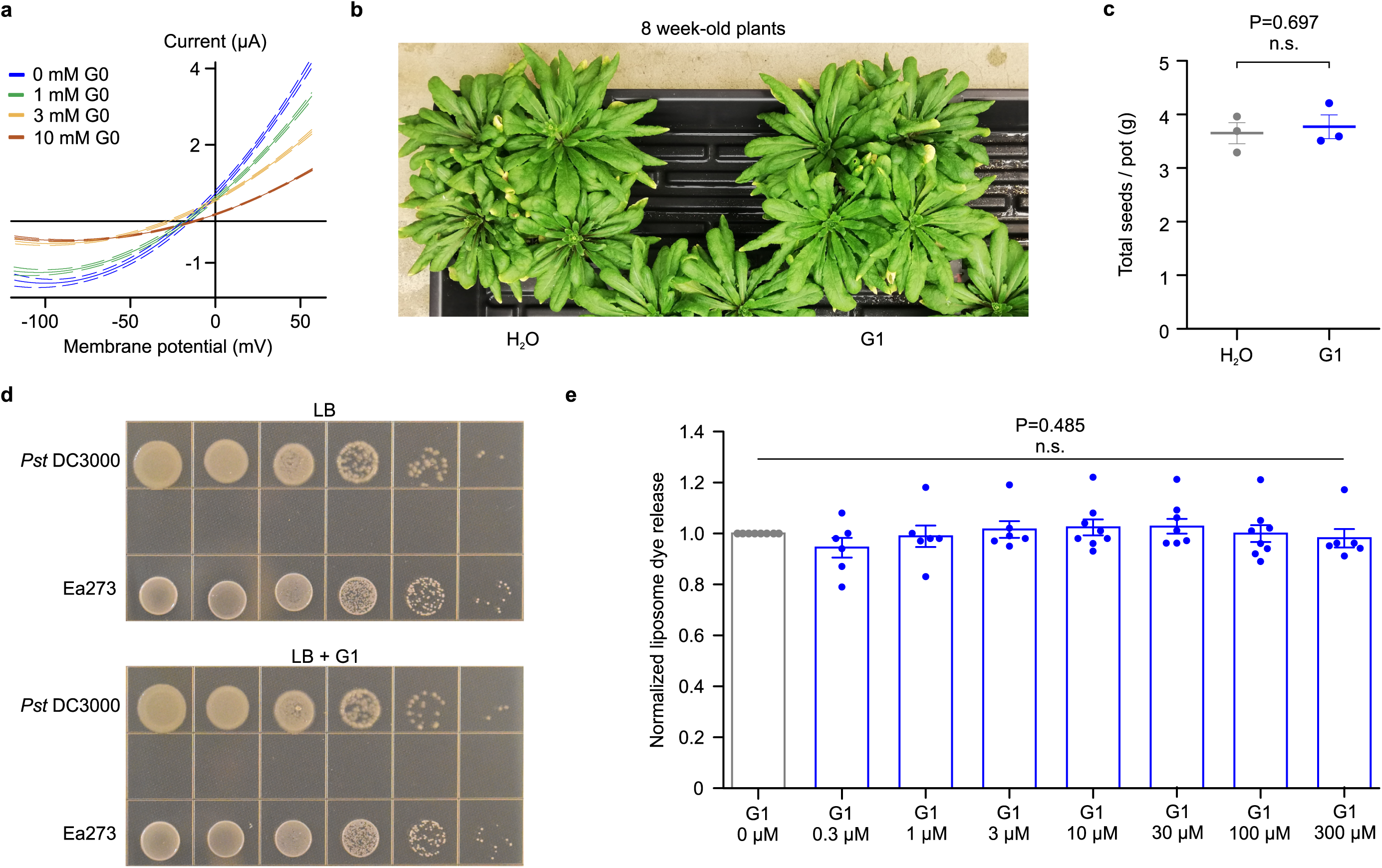
Effects of PAMAM inhibitors on DspE activities in oocytes and liposome and on plant phenotypes and bacterial growth. **a**, Inhibition of ion currents in oocytes. DspE-dependent currents were inhibited by G0 in a dose-dependent manner across all test pulses. Solid lines represent fit values across the entire voltage range, with dashed lines representing the lower and upper 95% confidence interval of a quadratic polynomial regression for each treatment after subtracting control values (n=5). **b**,**c**, PAMAM G1 does not affect plant growth or seed production of Arabidopsis Col-0 plant. Five-week-old Col-0 plants were sprayed with 10 µM PAMAM G1 every week until 12 weeks old. **b,** Picture of 8-week-old plants is shown here as an example. **c**, Seeds were collected at 15 weeks old. No obvious difference in plant appearance or seed production was observed. Data shown as mean ± SEM (n=3). **d**, PAMAM G1 does not impact bacteria multiplication *in vitro* (in LB agar medium). Ea273 or *Pst* DC3000 cells in the logarithmic growth phase were spotted, at a 10-fold serial dilution (from left to right), on low-salt LB agar plates with or without 50 μM PAMAM G1. Plates were kept at 28 for 2 days to visualize colonies. **e**, Normalized liposome dye release after addition of triton. Fluorescence readings of groups with increasing concentrations of G1 were normalized to the no compound controls and are shown as mean ± SEM (n=6 for 0.3, 1, 3, and 300 µM G1; n=8 for 10, 30, and 100 µM G1). Experiments were performed two (a) or three (b,c,d,e) times with similar results. Two-tailed student’s *t*-test (c), One-way ANOVA on Ranks (e), and regression (a) values are detailed in the Source Data files.

**Extended Data Figure 8.**
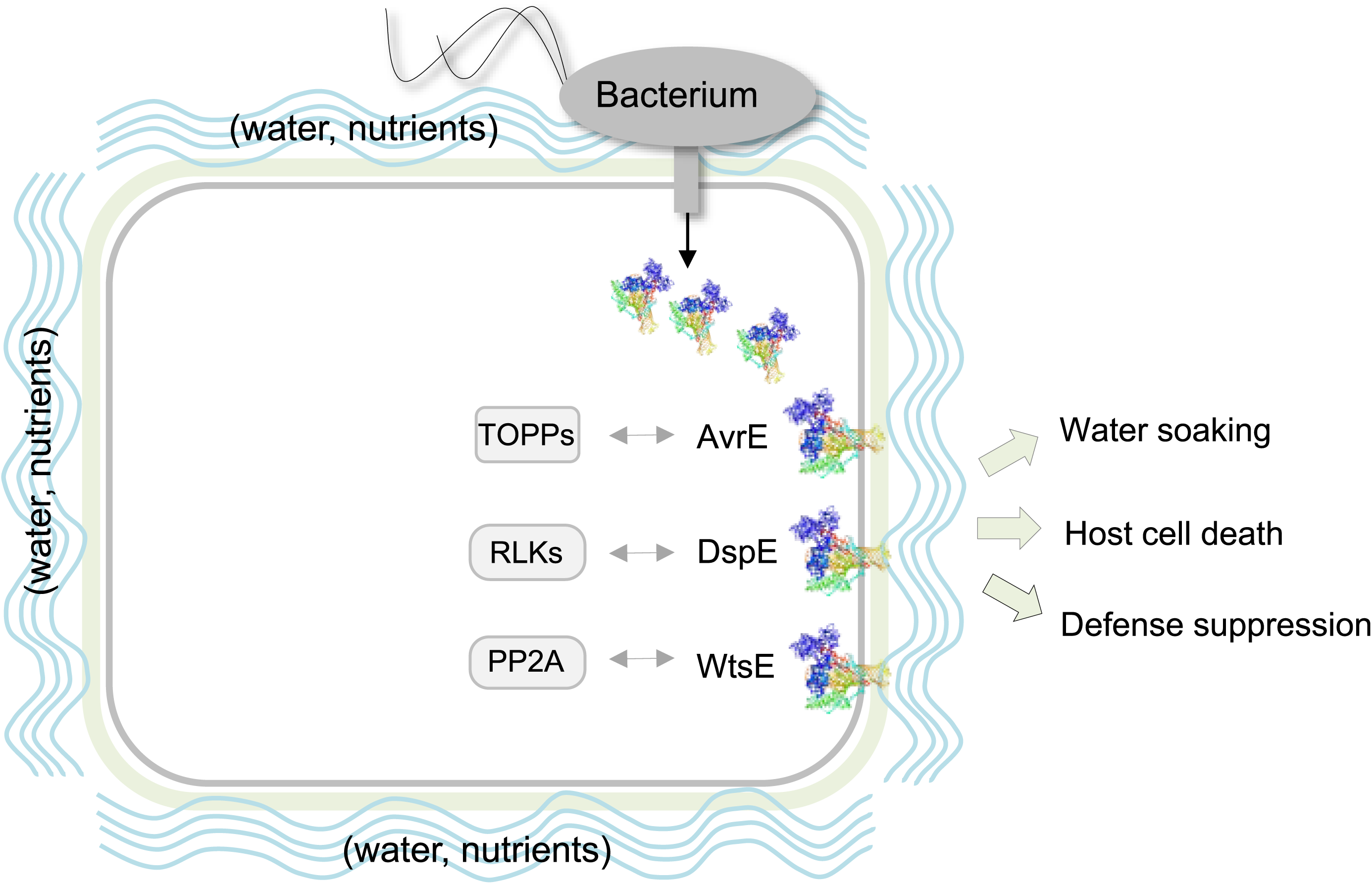
A working model for the molecular actions of AvrE/DspE-family effectors in plants. AvrE/DspE-family effectors act primarily as a novel class of water/solute-permeable channels dedicated to creating osmotic/water potential perturbation and a water/nutrient-rich apoplast, in which bacteria multiply within the infected plant tissues. AvrE/DspE-family effectors can additionally engage host proteins, including plant protein phosphatase PP2A subunits, type one protein phosphatases (TOPPs) and receptor-like kinases (RLKs), possibly to modulate AvrE/DspE-family channel properties or to optimize the pathogenic outcomes of AvrE/DspE-family channel activities, including water soaking, host cell death and defence suppression.

**Extended Data Figure 9.**
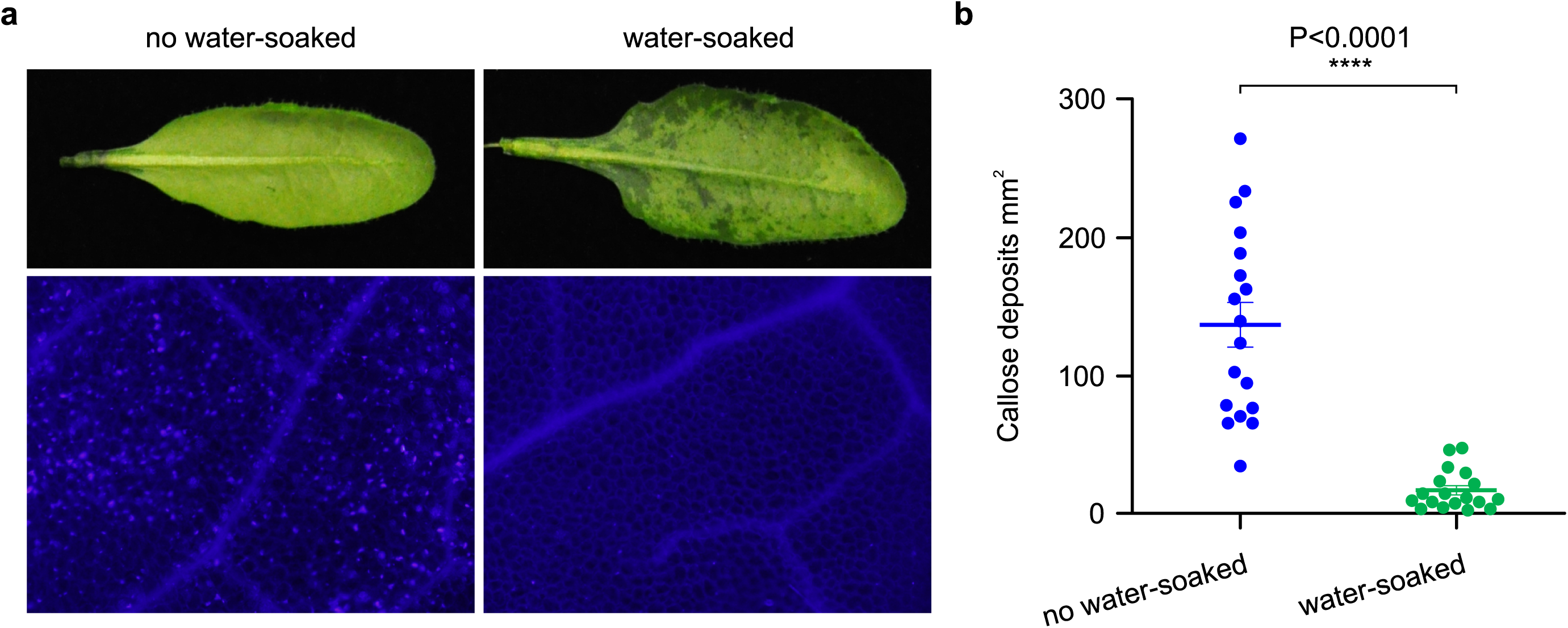
Water infiltration is sufficient to dampen callose deposition in Arabidopsis leaves. Arabidopsis Col-0 leaves were syringe-infiltrated with 1 µM flg22 and immediately covered by a clear plastic dome to maintain infiltrated leaves water-soaked (“water-soaked”; spotty, darker appearance) or air-dried to let infiltrated leaves returned to pre-infiltration appearance (∼1 h) and then covered by a clear plastic dome (“no water soaked”; uniform, lighter appearance). **a**, Leaf pictures and callose (bright dots) images were taken at 8 h post flg22 infiltration. **b**, Quantification of callose deposition (mean ± SEM; n=18). Experiments were performed three times with similar results. Two-tailed student’s *t*-test values are detailed in the Source Data files.

---

**Supplementary Figure 1.** Whole gel images for Extended Data Fig. 4a-f. Dotted boxes show image cropping.

**Supplementary Figure 2.** Whole gel images for Fig. 4c. Dotted boxes show image cropping.

**Supplementary Figure 3.** Whole gel images for Extended Data Fig. 4g,h. Dotted boxes show image cropping.

**Supplementary Video 1. AvrE rapid swelling and burst assay in *Xenopus* oocytes.** After 20 ng of cRNA was injected and time was allowed for the protein expression (24 h), oocytes were moved from a 200 mOsm saline solution to a 40 mOsm saline solution (5× dilution in ultrapure H_2_O). Note that AvrE-expressing oocytes further increase in size until burst mainly through the cRNA injection site where the extracellular matrix layer surrounding the plasma membrane is weaker than the rest of the oocyte. Cells were imaged every 20s under a stereomicroscope (7.5× magnification), pictures were assembled in order, and the final time-lapse video is at 3.33Hz. Video was assembled in iMovie v10.3.4, Apple Inc.

**Supplementary Video 2. DspE rapid swelling and burst assay in *Xenopus* oocytes**. After 2 ng of cRNA was injected and time was allowed for the protein expression (24 h), oocytes were moved from a 200 mOsm saline solution to a 40 mOsm saline solution (5× dilution in ultrapure H_2_O). Note that DspE-expressing oocytes further increase in size until burst mainly through the cRNA injection site where the extracellular matrix layer surrounding the plasma membrane is weaker than the rest of the oocyte. Cells were imaged every 20s under a stereomicroscope (7.5× magnification), pictures were assembled in order, and the final time-lapse video is at 3.33Hz. Video was assembled in iMovie v10.3.4, Apple Inc.

## Notes

### Competing Interest Statement

The authors have declared no competing interest.

