## Supplementary figures and images for "Bacterial pathogens deliver water/solute-permeable channels as a virulence strategy"

### Supplementary Figure 1

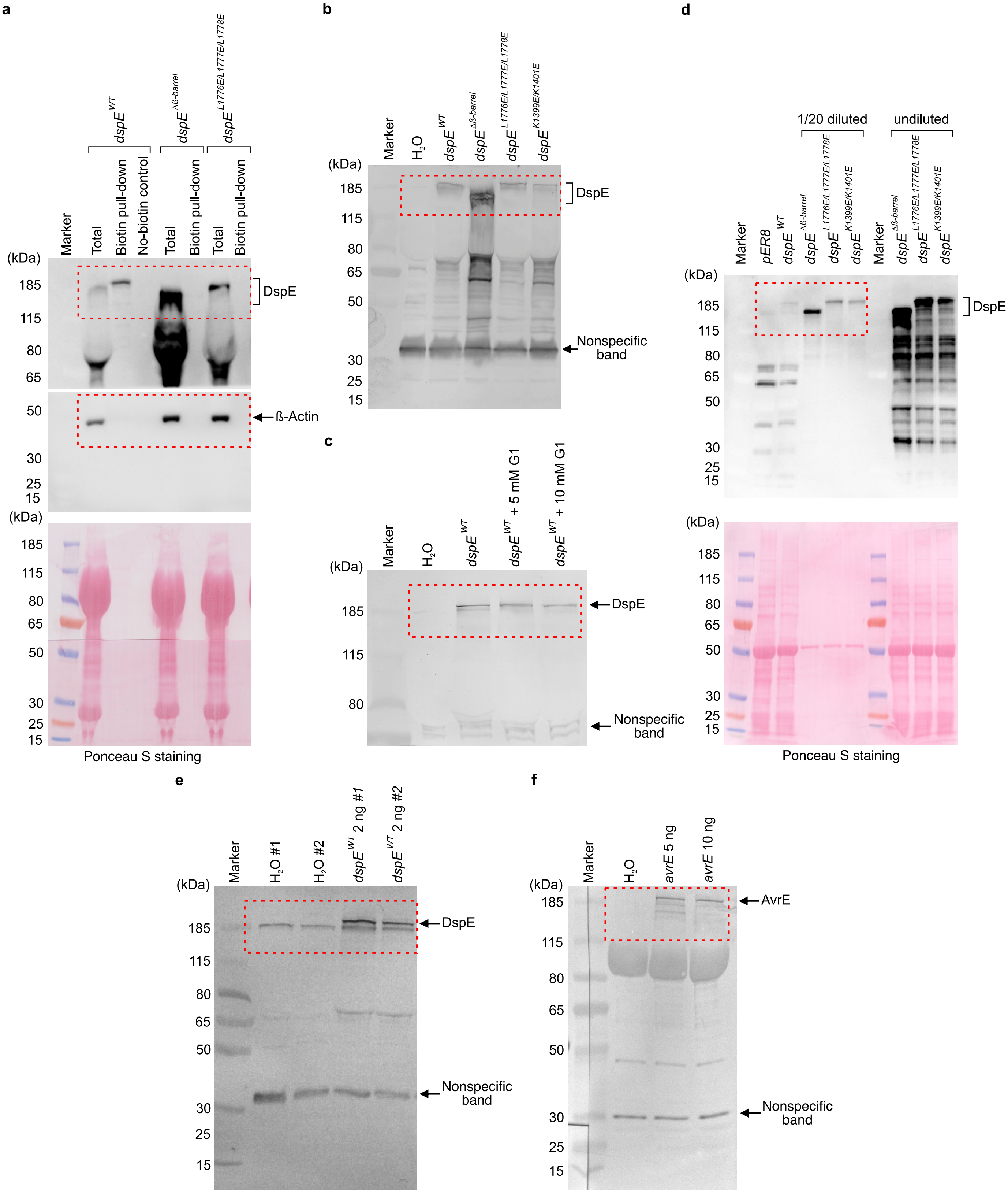

### Supplementary Figure 2

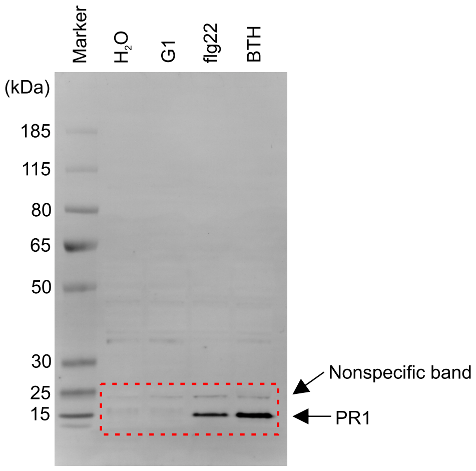

### Supplementary Figure 3

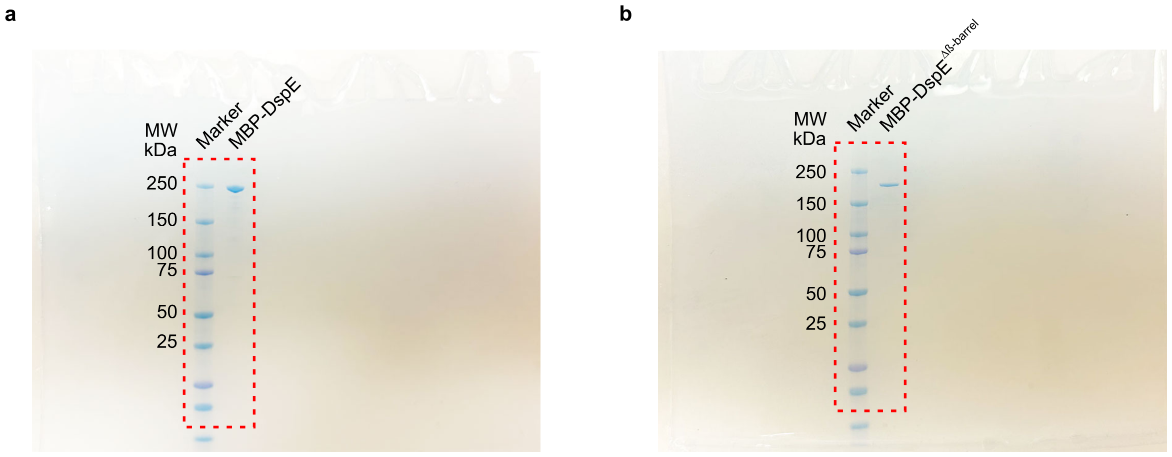
